# Tricalbin proteins regulate plasma membrane phospholipid homeostasis

**DOI:** 10.1101/2021.06.04.447076

**Authors:** Ffion B. Thomas, Deike J. Omnus, Jakob M. Bader, Gary H. C. Chung, Nozomu Kono, Christopher J. Stefan

## Abstract

The evolutionarily conserved extended synaptotagmin (E-Syt) proteins are calcium-activated lipid transfer proteins that function at contacts between the endoplasmic reticulum and plasma membrane (ER-PM contacts). However, roles of the E-Syt family members in PM lipid organisation remain unclear. Among the E-Syt family, the yeast tricalbin (Tcb) proteins are essential for PM integrity upon heat stress, but it is not known how they contribute to PM maintenance. Using quantitative lipidomics and microscopy, we find that the Tcb proteins regulate phosphatidylserine homeostasis at the PM. Moreover, upon heat-induced membrane stress, Tcb3 co-localises with the PM protein Sfk1 that is implicated in PM phospholipid asymmetry and integrity. The Tcb proteins also promote the recruitment of Pkh1, a stress-activated protein kinase required for PM integrity. Phosphatidylserine has evolutionarily conserved roles in PM organisation, integrity, and repair. We suggest that phospholipid regulation is an ancient essential function of E-Syt family members in PM integrity.

## Introduction

Maintaining the mechano-chemical properties of the plasma membrane (PM) is essential to vital processes including selective ion and nutrient transport, as well as size and shape control in all living cells. Accordingly, the PM has a distinctive lipid composition in eukaryotic cells, including high sterol and sphingolipid content as well as an enrichment of phosphatidylserine (PS) in its cytosolic leaflet that endows the PM with its unique identity, biophysical properties, organisation, and integrity (Schneiter et al., 1999; van Meer et al., 2008; Yeung et al., 2008; Lingwood and Simons, 2010; Bigay and Antonny, 2012; Holthuis and Menon, 2014). PM lipid composition is achieved and maintained, as needed, through the selective delivery of lipids from the endoplasmic reticulum (ER) where they are synthesized to the PM by vesicular and non-vesicular transport pathways. Vesicular lipid trafficking occurs via the secretory pathway alongside PM-bound proteins (Klemm et al., 2009; Fairn et al., 2011). It is also clear that lipid transfer proteins mediate non-vesicular lipid exchange between the ER and PM in the control of PM lipid composition and homeostasis (Kaplan and Simoni, 1985; Urbani and Simoni, 1990; Vance et al., 1991; Holthuis and Menon, 2014; Wong et al., 2019). Membrane contact sites between the ER and the PM, termed ER-PM contacts, are proposed to serve as integral sites for the coordinated regulation of lipid metabolism and transport (Pichler et al., 2001; Chang et al., 2017; Balla et al., 2020; Nishimura and Stefan, 2020). However, a strict requirement for ER-PM contacts in non-vesicular transport of lipids from the ER to PM has been recently questioned (Quon et al., 2018; Wang et al., 2020), necessitating further evaluation of the vital roles of ER-PM contacts, as well as the proteins proposed to form and function at these important cellular structures.

Whilst roles of several tether and lipid transfer proteins identified at ER-PM contacts are established, functions of the extended synaptotagmin (E-Syt) protein family members are incompletely understood and even controversial. The E-Syt proteins, as well as their budding yeast orthologs, named tricalbins, are anchored in the ER membrane via a N-terminal hairpin (Giordano et al., 2013) and interact with the PM in a phosphoinositide lipid- and Ca^2+^-dependent manner via their multiple C-terminal cytoplasmic C2 domains (Chang et al., 2013; Giordano et al., 2013; Idevall-Hagren et al., 2015; Saheki et al., 2016; Bian et al., 2018). They also feature a central cytosolic synaptotagmin-like, mitochondrial (SMP) domain that dimerizes and contains a deep hydrophobic groove previously shown to bind and transport lipids *in vitro* (Schauder et al., 2014; Saheki et al., 2016; Yu et al., 2016; Bian et al., 2018; Bian and De Camilli, 2019; Qian et al., 2021). Whilst cellular roles of E-Syt proteins as ER-PM tethers are well-described (Giordano et al., 2013, Fernández-Busnadiego et al. 2015), precise roles of the E-Syts the control of membrane lipid dynamics *in vivo* are enigmatic. One study demonstrated a role of the E-Syts in the transfer of diacylglycerol from the PM to the ER during the phosphoinositide cycle, but loss of the E-Syt1/2/3 proteins had no significant effect on phosphoinositide lipid synthesis or the homeostasis of other phospholipids at the PM (Saheki et al., 2016). An earlier study found that depletion of the E-Syt1/2 proteins impaired the re-synthesis of phosphatidylinositol (4,5)-bisphosphate, commonly termed PI(4,5)P_2_, during the phosphoinositide cycle and suggested a role of E-Syt-mediated ER-PM contacts in the transfer of phosphatidylinositol from the ER to the PM (Chang et al., 2013). Yet another study has even suggested a role for E-Syt2 in PI(4,5)P_2_ turnover (Dickson et al., 2016). Whilst the findings in these studies are not necessarily mutually exclusive or contradictory, they highlight unresolved issues regarding the roles of the E-Syts in PM lipid homeostasis.

The budding yeast E-Syt orthologs, the tricalbins (Tcb1/2/3), have also been shown to play a role in ER-PM contact formation (Manford et al., 2012; Toulmay and Prinz, 2012; Collado et al., 2019; Hoffmann et al., 2019). Two recent studies used advanced cryo-electron tomography to reveal peaks of extreme ER membrane curvature at Tcb-mediated ER-PM contacts (Collado et al., 2019; Hoffmann et al., 2019). In particular, one study also found that Tcb-dependent ER-PM contacts are induced upon PM stress conditions and are required to maintain PM integrity upon stress conditions (Collado et al., 2019). Moreover, computational modelling suggested that the regions of extreme ER membrane curvature may facilitate lipid transfer from the ER to the PM at Tcb-mediated ER-PM contacts (Collado et al., 2019). However, potential functions of the Tcb proteins in lipid transport at ER-PM contacts remain to be experimentally tested. Consequently, mechanistic insight into the roles of the Tcb proteins in PM lipid homeostasis and integrity is lacking.

In this study, we use quantitative lipidomics and microscopy approaches to elucidate roles of the Tcb proteins in the control of PM lipid composition. The data show that the Tcb proteins regulate phospholipid homeostasis at the PM upon stress conditions. In particular, the results are consistent with a role of ER-PM contacts in the delivery of certain PS and phosphatidylethanolamine (PE) species, but not phosphatidylinositol (PI), from the ER to the PM. Furthermore, we find that Tcb3 co-localises with the PM protein Sfk1, an ortholog of the mammalian TMEM150/FRAG1/DRAM proteins, that is implicated in stress-induced PI(4,5)P_2_ synthesis and phospholipid asymmetry at the PM (Audhya and Emr, 2002; Chung et al., 2015; Mioka et al., 2018). Finally, we find that the Tcb proteins promote the recruitment of Pkh1, a stress-activated protein kinase required for PM integrity, to the PM upon stress conditions. Altogether, our findings indicate that the Tcb proteins function as inducible ER-PM tethers necessary for PM phospholipid homeostasis upon stress conditions, providing new mechanistic insight into Tcb protein function at ER-PM contact sites.

## Results

### The tricalbins regulate phosphoinositide homeostasis at the PM

Several proteins have been shown to form and function at ER-PM contacts in yeast, including Scs2/22 (VAP orthologs), Ist2 (ANO8/TMEM16 ortholog), Lam1-4 (GRAMD1/Aster orthologs), and the tricalbin (Tcb) proteins (Loewen et al., 2007; Manford et al., 2012; Gatta et al., 2015) (**Figure 1A**). The aim of this study is to elucidate functions of the Tcb proteins at ER-PM contacts that have remained incompletely understood. Under normal growth conditions, loss of Tcb1, Tcb2, and Tcb3 (in *tcb1/2/3*Δ cells) does not result in obvious effects on ER-PM tethering (Manford et al., 2012) or phospholipid homeostasis (**Figures 2** and **3**). However, roles of the Tcb proteins in ER-PM tethering become apparent upon loss of additional proteins that form ER-PM contacts, including Scs2/22 and Ist2 (**Figure 1A**) (Manford et al., 2012; Collado et al., 2019). Specifically, Tcb-mediated ER-PM contacts are induced upon loss of the Scs2/22 and Ist2 proteins (in *scs2/22*Δ *ist2*Δ triple mutant cells) (Collado et al., 2019), and loss of Tcb1/2/3 in combination with loss of Scs2/22 and Ist2 results in additive defects in ER-PM tethering (Manford et al., 2012). However, specific roles of the Tcb proteins in PM lipid homeostasis have not been rigorously examined. We therefore monitored the consequential effects of loss of the Tcb proteins in *scs2/22*Δ *ist2*Δ triple mutant cells on PM lipid homeostasis.

**Figure 1.**
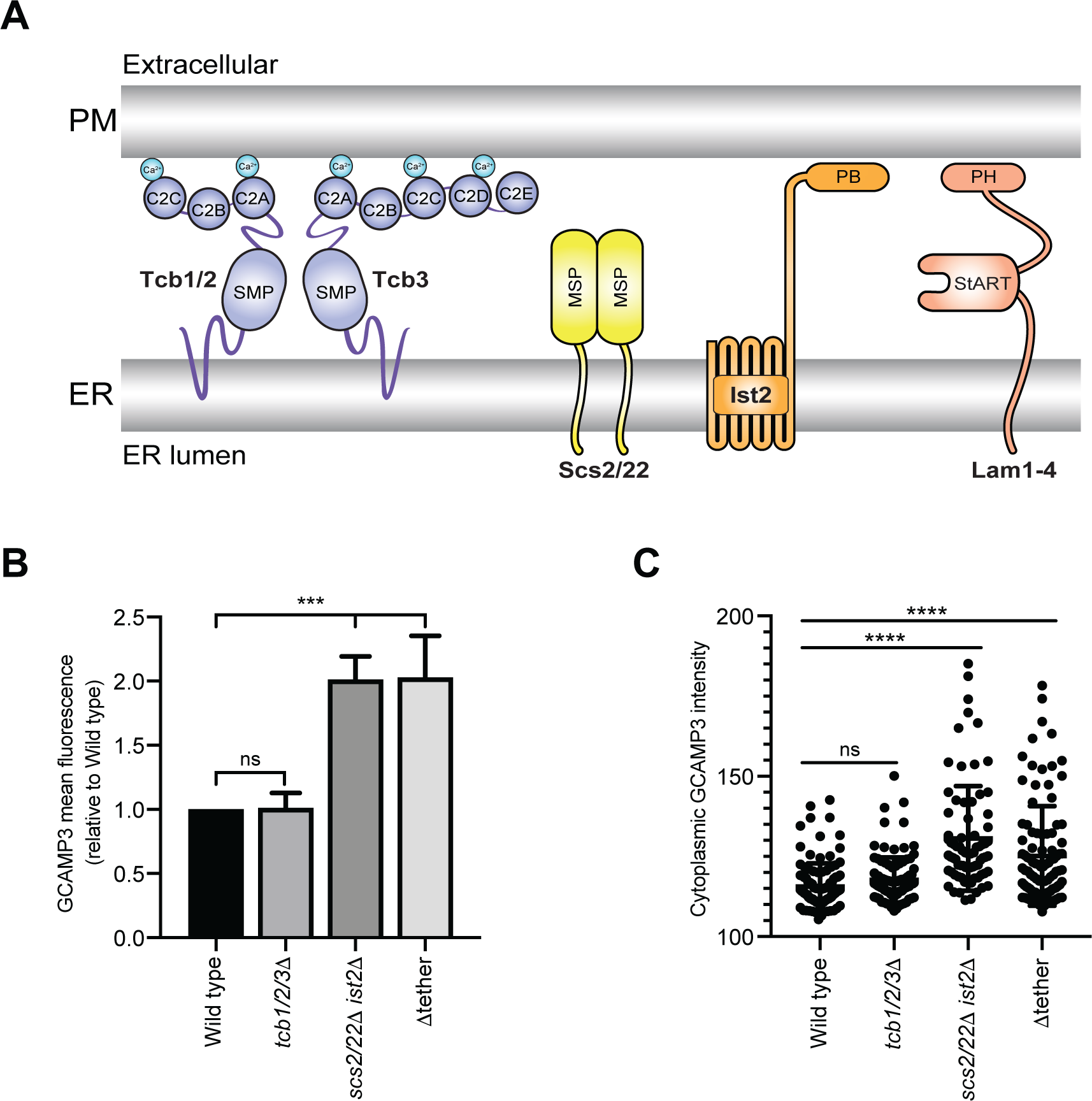
Cytoplasmic calcium levels are increased in *scs2/22*Δ *ist2*Δ and Δtether mutant cells. (**A**) Schematic representations of ER-PM tethering proteins found in budding yeast. These include the tricalbins (Tcb), Scs2 and Scs22, Ist2 and Lam proteins. This study is focused on elucidating specific roles of the Tcb proteins in PM homeostasis. Abbreviations: SMP, Synaptotagmin-like mitochondrial-lipid-binding domain; C2, C2 domain; MSP, Major sperm protein domain; PB, Polybasic stretch, StART, StAR-related lipid-transfer domain; PH, Pleckstrin homology domain. (**B**) Mean fluorescence intensity of cytoplasmic GCaMP3 reporter of wild type, *tcb1/2/3*Δ, *scs2/22*Δ *ist2*Δ and Δtether cells as measured by flow cytometry (100,000 cells measured per experiment). Data represents mean ± standard deviation from three independent experiments. *** p > 0.001. (**C**) Quantitation of GCaMP3 signal in the cytosol of wild type, *tcb1/2/3*Δ, *scs2/22*Δ *ist2*Δ and Δtether cells measured by fluorescence microscopy. Data represents mean ± standard deviation. Total number of cells analysed in three independent experiments: wild type n=109, *tcb1/2/3*Δ n=103, *scs2/22*Δ *ist2*Δ n=97, Δtether n=100. **** p > 0.0001. Also see **Figure S1**.

**Figure 2.**
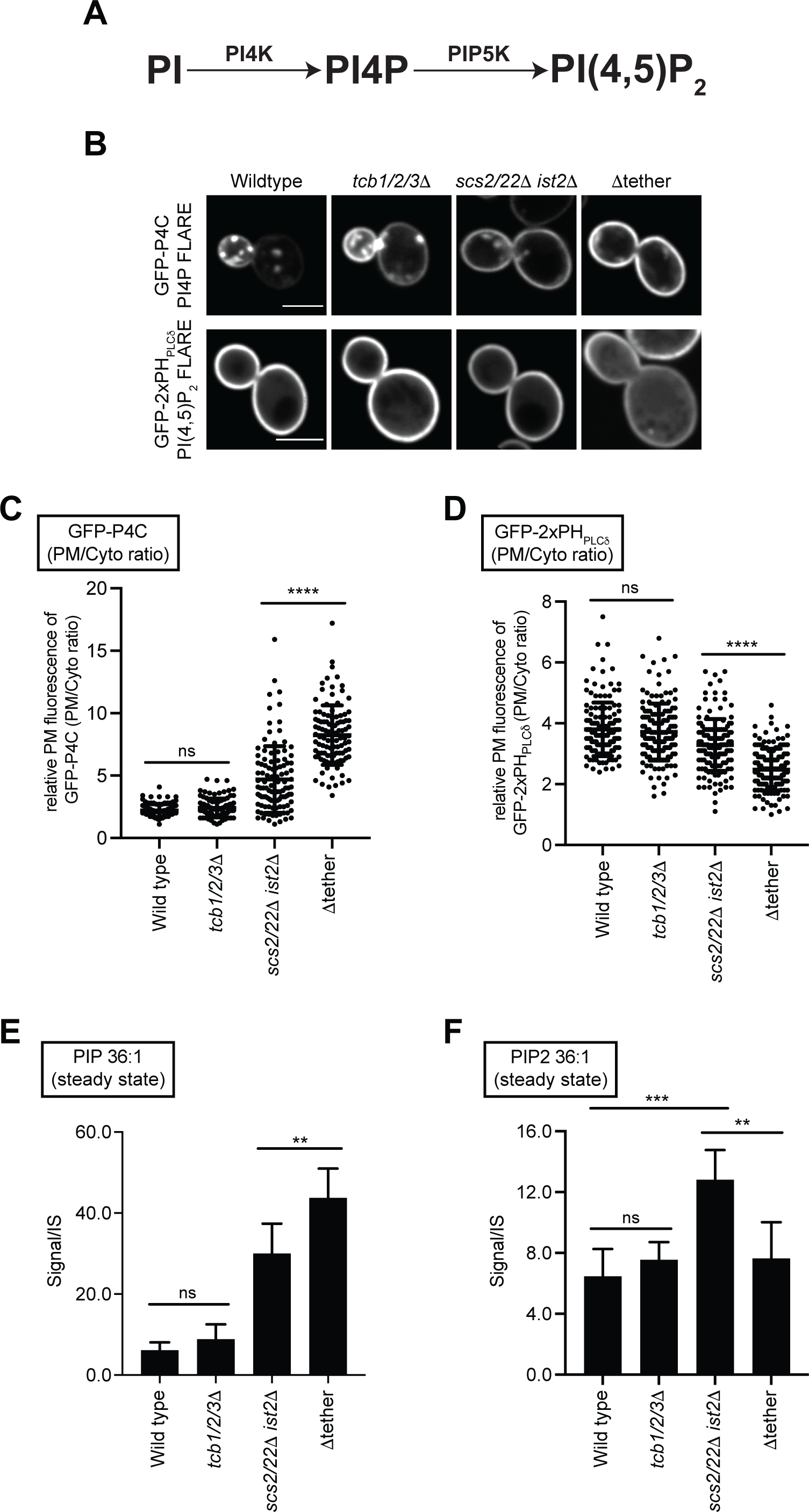
Tricalbin proteins regulate phosphoinositide homeostasis at the PM. (**A**) Schematic representation of PI4P and PI(4,5)_2_ production at the PM and the kinases involved. (**B**) PI4P (GFP-P4C) and PI(4,5)P_2_ (GFP-2xPH_PLCδ_) FLARE localisation in wild type, *tcb1/2/3*Δ, *scs2/22*Δ *ist2*Δ and Δtether cells. Scale bars, 4 μm. (**C**) Quantitation of GFP-P4C intensity at the PM of the mother cell. Data represents mean ± standard deviation. Total number of cells analysed in three independent experiments: wild type n=104, *tcb1/2/3*Δ n=102, *scs2/22*Δ *ist2*Δ n=107, Δtether n=105. **** p > 0.0001. (**D**) Quantitation of GFP-2xPH_PLCδ_ intensity at the PM of the mother cell. Data represents mean ± standard deviation. Total number of cells analysed in three independent experiments: wild type n=146, *tcb1/2/3*Δ n=149, *scs2/22*Δ *ist2*Δ n=146, Δtether n=147). **** p > 0.0001. (**E** and **F**) Lipidomic analysis of PIP 36:1 and PIP2 36:1 species in wild type, *tcb1/2/3*Δ, *scs2/22*Δ *ist2*Δ and Δtether cells. Data represents mean ± standard deviation (N=5). ** p > 0.01, *** p > 0.001. Also see **Figure S2**.

**Figure 3.**
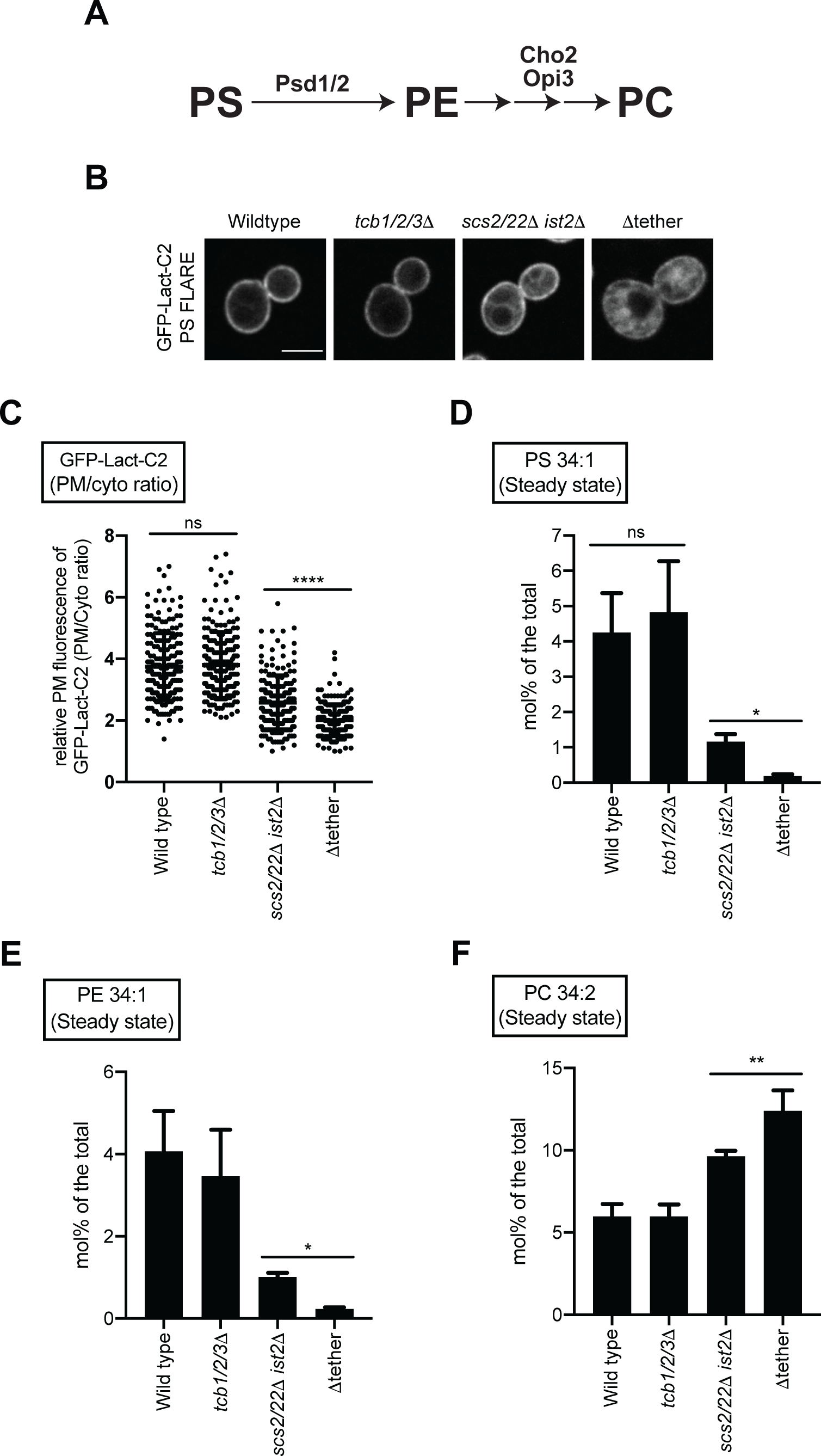
Tricalbin proteins regulate phospholipid homeostasis at the PM. (**A**) Schematic representation of PS, PE, and PC production and the enzymes involved. (**B**) PS FLARE (GFP-Lact-C2) localisation in wild type, *tcb1/2/3*Δ, *scs2/22*Δ *ist2*Δ and Δtether cells. Scale bars, 4 μm. (**C**) Quantitation of relative GFP-Lact-C2 intensity at the PM of the mother cell. Data represents mean ± standard deviation. Total number of cells analysed in three independent experiments: wild type n=196, *tcb1/2/3*Δ n=194, *scs2/22*Δ *ist2*Δ n=187, Δtether n=184. **** p > 0.0001. (**D**, **E,** and **F**) Lipidomic analyses of PS 34:1, PE 34:1 and PC 34:2 species in wild type, *tcb1/2/3*Δ, *scs2/22*Δ *ist2*Δ and Δtether cells. Data represents mean ± standard deviation (N=3). * p > 0.1, ** p > 0.01. Also see **Figure S3**.

The E-Syt1 protein has been shown to be specifically activated upon increases cytoplasmic calcium (Ca^2+^) (Fernandez-Busnadiego et al., 2015; Idevall-Hagren et al., 2015; Bian et al., 2018). To confirm that Tcb-mediated ER-PM contacts are induced in cells lacking Scs2/22 and Ist2 (*scs2/22*Δ *ist2*Δ cells) (Collado et al., 2019), we performed control experiments to monitor the cytoplasmic Ca^2+^ reporter GCaMP3 in *scs2/22*Δ *ist2*Δ cells. Consistent with Tcb activation, GCaMP3 fluorescence was increased in *scs2/22*Δ *ist2*Δ cells as compared to wild-type cells, as measured by high-content quantitative flow cytometry assays (2-fold; **Figure 1B**) and fluorescence microscopy (**Figure 1C**). Furthermore, impairment of Ca^2+^ influx in the *scs2/22*Δ *ist2*Δ mutant cells phenocopied loss of the Tcb proteins in this mutant background (*scs2/22*Δ *ist2*Δ *tcb1/2/3*Δ cells, also named Δtether cells). Specifically, loss of the stretch-activated Ca^2+^ channel Mid1 (Iida et al., 1994) in the *scs2/22*Δ *ist2*Δ mutant cells (*scs2/22*Δ *ist2*Δ *mid1*Δ) conferred increased resistance to the drug myriocin, similar to the Δtether mutant cells (**Figure S1A**). Thus, cytoplasmic Ca^2+^ is elevated in *scs2/22*Δ *ist2*Δ cells, consistent with a previous study (Kato et al., 2017), and the Mid1-dependent high-affinity Ca^2+^ influx system (termed HACS) (Muller et al., 2001) may contribute to Tcb protein function.

Following these control experiments, we investigated roles of the Tcb proteins in phosphoinositide metabolism at the PM using quantitative microscopy and mass spectrometry-based lipidomics. PI(4,5)P_2_ and its precursor PI4P (phosphatidylinositol 4-phosphate) are the two major phosphoinositide species present at the PM in yeast and are generated through sequential phosphorylation of PI at the PM (**Figure 2A** and **S2A**). As expected, localisation of the PI4P and PI(4,5)P_2_ biosensor FLAREs (fluorescent lipid-associated reporters), GFP-P4C and GFP-2xPH_PLCδ_ respectively, was not significantly affected by loss of the Tcb proteins (*tcb1/2/3*Δ cells), as compared to wild type control cells (**Figures 2B, C** and **D**). In contrast, there was a significant increase in the localisation of the PI4P FLARE (GFP-P4C) at the PM in *scs2/22*Δ *ist2*Δ cells, as compared to wild type control cells (**Figures 2B** and **C**). Previous studies have shown that Scs2/22 and Ist2 recruit the PI4P exchange proteins Osh2, Osh3, Osh6, and Osh7 to ER-PM contacts (Loewen and Levine, 2005; D’Ambrosio et al., 2020) and loss of Scs2/22 and Ist2 results in increased PI4P levels (Manford et al., 2012). As Tcb-mediated ER-PM contacts are induced in cells lacking Scs2/22 and Ist2 (*scs2/22*Δ *ist2*Δ), we next monitored distribution of the PI4P and PI(4,5)P_2_ FLAREs in *scs2/22*Δ *ist2*Δ cells that also lacked the Tcb proteins (Δtether). Interestingly, whilst there was a significant increase in the PI4P FLARE at the PM of Δtether cells compared to *scs2/22*Δ *ist2*Δ cells (**Figures 2B** and **C**), there was a decrease in the relative levels of the PI(4,5)P_2_ FLARE at the PM in Δtether cells compared to *scs2/22*Δ *ist2*Δ cells (**Figures 2B** and **D**). Thus, the Tcb proteins are not required for PI4P generation at the PM, but they may contribute to PI(4,5)P_2_ homeostasis at the PM.

Next, we confirmed the PI4P and PI(4,5)P_2_ FLARE results using quantitative lipidomics. For these experiments, we measured levels of phosphoinositide species in each of the ER-PM tether mutants by liquid chromatography-electrospray ionization-tandem mass spectrometry (LC-ESI-MS/MS) analysis (Clark et al., 2011). Consistent with the microscopy results, levels of mono-unsaturated phosphatidylinositol phosphate (PIP), that constitute the majority of PI4P species generated at the PM (Wenk et al., 2003; Nishimura et al., 2019), progressively increased between wild type control cells, *scs2/22*Δ *ist2*Δ cells, and Δtether cells with the Δtether cells containing significantly higher PIP steady-state levels than the *scs2/22*Δ *ist2*Δ cells (**Figures 2E** and **S2B**). Remarkably, however, the level of mono-unsaturated phosphatidylinositol bis-phosphate (PIP_2_) did not reflect the increase in PIP steady-state levels in the Δtether cells (**Figures 2F** and **S2B**). In particular, *scs2/22*Δ *ist2*Δ cells displayed increases in both 36:1 PIP and 36:1 PIP_2_ steady-state levels compared to wild type control cells (5-fold and 2-fold, respectively; **Figures 2E** and **2F**). However, whilst the Δtether cells (also lacking the Tcb proteins) displayed a further increase in 36:1 PIP levels (1.5-fold as compared to *scs2/22*Δ *ist2*Δ cells; **Figure 2E**), 36:1 PIP_2_ steady-state levels were decreased (1.7-fold as compared to *scs2/22*Δ *ist2*Δ cells; **Figure 2F**). Likewise, 34:1 PIP levels were increased in the Δtether cells as compared to *scs2/22*Δ *ist2*Δ cells (1.8-fold; **Figure S2B**), whereas 34:1 PIP_2_ levels were slightly decreased in the Δtether cells (**Figure S2B**). As expected, no significant differences were detected between *tcb1/2/3*Δ cells and wild type control cells (**Figure 2** and **S2B**). Intriguingly, none of the strains tested showed significant changes in steady-state levels of any PI species (**Figure S2B**). Previous studies have implicated E-Syt family members in recycling diacylglycerol (DAG) from the PM to the ER and in the transfer of PI from the ER to the PM during the phosphoinositide cycle (Chang et al., 2013; Saheki et al., 2016; Nath et al., 2020). However, whilst PI homeostasis and PI4P synthesis at the PM apparently do not require the Tcb proteins, conversion of PI4P to PI(4,5)P_2_ at the PM is impaired. Altogether, these results point to a function that has yet to be described for metazoan E-Syt family members.

### The tricalbins regulate phospholipid homeostasis at the PM

We next investigated how the Tcb proteins might contribute to PI(4,5)P_2_ homeostasis at the PM. PI4P 5-kinase (PIP5K) activity has been shown to be directly regulated by anionic phospholipids, including PS that is enriched in the cytosolic leaflet of the PM and contributes to its overall negative charge (Fairn et al., 2009; Nishimura et al., 2019). PS is synthesized in the ER via the CDP-DAG pathway in yeast and either converted to other phospholipids (**Figures 3A** and **S3A**) or transferred to the PM via vesicular and non-vesicular transport pathways. The generation of Tcb-mediated ER peaks facing the PM has been suggested to facilitate non-vesicular lipid transport from the ER to the PM (Collado et al., 2019). We therefore considered whether the Tcb proteins regulate non-vesicular delivery of PS to the PM. First, we analysed the distribution of a PS FLARE (GFP-Lact-C2) in the same series of ER-PM tether mutants by quantitative microscopy. Under normal growth conditions, loss of the Tcb proteins did not significantly affect the localisation of GFP-Lact-C2 at the PM as compared to wild type control cells (**Figures 3B** and **C**). However, the relative fluorescence intensity of GFP-Lact-C2 at the PM was significantly reduced in Δtether cells as compared to *scs2/22*Δ *ist2*Δ cells (**Figures 3B** and **C**). This effect was also observed when comparing the *scs2/22*Δ *ist2*Δ and *scs2/22*Δ *ist2*Δ *mid1*Δ mutants (**Figure S1B**).

We further investigated roles of the Tcb proteins in membrane lipid homeostasis using quantitative lipidomics. Consistent with the microscopy results, steady-state levels of PS were reduced in the *scs2/22*Δ *ist2*Δ and Δtether mutant cells, but not in *tcb1/2/3*Δ cells, as compared to wild type control cells (**Figures 3D** and **S3B**). Moreover, species-level analyses revealed a specific reduction in mono-unsaturated 34:1 PS, which is the major PS species enriched at the PM (Schneiter et al., 1999), in Δtether cells compared to *scs2/22*Δ *ist2*Δ cells (>7-fold; **Figures 3D** and **S3B**). In contrast, there were no significant differences in the levels of di-unsaturated 32:2 or 34:2 PS in any of the strains examined (**Figure S3B**), indicating that PS synthesis *per se* was not completely disrupted by loss of the ER-PM tether proteins. Consistent with this, *cho1*Δ mutant cells defective in PS synthesis rely upon exogenous ethanolamine or choline in order to support PE and PC synthesis for growth (Atkinson et al., 1980), but the Δtether cells are viable in the absence of exogenous ethanolamine and choline and grow on standard media (see **Figure S1**). Thus, whilst levels of ‘ER-like’ di-unsaturated PS species are retained, loss of Tcb activity in the *scs2/22*Δ *ist2*Δ cells resulted in a decrease in PM-specific mono-unsaturated 34:1 PS species, consistent with impaired transport of newly synthesized mono-unsaturated 34:1 PS from the ER to the PM.

In yeast, PS can be converted to phosphatidylethanolamine (PE) through decarboxylation reactions carried out by Psd1 and Psd2, and PE can be methylated by Cho2 and Opi3 to generate phosphatidylcholine (PC) (**Figures 3A** and **S3A**). A previous study found that loss of Psd1 is synthetic lethal in the Δtether cells (Wang et al., 2020), suggesting that PS species may be converted to other phospholipids in the Δtether cells in order to prevent membrane organelle dysfunction. Accordingly, quantitative lipidomic analyses in this study revealed specific changes in PE and PC species in the mutant cells. First, there were significant decreases in mono-unsaturated 32:1 and 34:1 PE, the major PE species enriched at the PM (Schneiter et al., 1999), in the *scs2/22*Δ *ist2*Δ and Δtether mutant cells as compared to wild type cells (**Figures 3E** and **S3B**). Second, mono-unsaturated 34:1 PE levels were further decreased in the Δtether cells as compared to *scs2/22*Δ *ist2*Δ cells (4-fold; **Figures 3E** and **S3B**), revealing a role of the Tcbs in regulation of 34:1 PE homeostasis. In contrast, there was no significant difference in the levels of di-unsaturated 34:2 PE in any of the strains examined (**Figure S3B**), indicating that PE synthesis was not blocked in the ER-PM tether mutants. Furthermore, there was a significant increase in di-unsaturated forms of PC (32:2, 34:2, and 36:2 PC) in the *scs2/22*Δ *ist2*Δ and Δtether mutant cells as compared to wild type cells (**Figures 3F** and **S3B**). Finally, di-unsaturated PC species were further increased in the Δtether cells as compared to *scs2/22*Δ *ist2*Δ cells (**Figures 3F** and **S3B**). Altogether, the results suggest that the Tcb proteins are needed to sustain delivery of 34:1 PS (and 34:1 PE) to the PM in the absence of Scs2/22 and Ist2. Furthermore, mono-unsaturated PS (and PE) species retained in the ER may be further desaturated and converted to di-unsaturated PC species in the mutant cells. Although, the data do not exclude a possible role of the Kennedy pathway in the accumulation of PC species in the *scs2/22*Δ *ist2*Δ and Δtether mutants.

### Tricalbins are required for PM phospholipid homeostasis and PM integrity under heat shock conditions

Sfk1 is an integral PM protein that has been implicated in PM phospholipid asymmetry and PM integrity (Mioka et al., 2018), as well as heat-induced PI(4,5)P_2_ synthesis (Audhya and Emr, 2002). Likewise, mammalian Sfk1 orthologs, the TMEM150 proteins, have been implicated in PI(4,5)P_2_ re-synthesis following phospholipase C activation (Chung et al., 2015). We observed that the ER-localised Tcb3 protein is in close proximity to the integral PM protein Sfk1, as assessed by bimolecular fluorescence complementation (BiFC) split GFP assays (**Figures 4A** and **B**). In control experiments, Tcb3 split GFP fusion proteins formed cortical patches dependent on the Tcb3 C2 domains (**Figure S4A**) that target Tcb3 to the cortical ER (Manford et al., 2012). In addition, Sfk1 and Tcb3 did not associate with the ER membrane protein Sec61 in BiFC split GFP assays (**Figure S4A**), indicating a specific association between Sfk1 and Tcb3. Consistent with this, the C-terminal cytoplasmic domain of Sfk1 was required for efficient association with Tcb3, as the BiFC split GFP signal intensity significantly decreased in cells expressing a truncated version of Sfk1 (Sfk1^Δ286-353^) (**Figure 4B**). As mentioned, Sfk1 has been implicated in PM lipid organisation and integrity (Mioka et al., 2018), as well as heat-induced PI(4,5)P_2_ synthesis at the PM (Audhya and Emr, 2002). Tcb3 has been shown to be necessary for the formation of heat-induced ER-PM contacts and for the maintenance of PM integrity upon heat stress conditions (Collado et al., 2019). We therefore assessed the localisation of Tcb3-GFP with Sfk1-mCherry following a brief heat shock (10 min 42°C). Sfk1 and Tcb3 displayed increased co-localisation at the cell cortex upon a brief incubation at 42°C (**Figures 4C** and **D**). In control experiments, a shift to 42°C induced heterogeneous transient cytoplasmic Ca^2+^ bursts up to 15-fold higher than basal levels at 26°C, as detected by GCaMP3 fluorescence (**Figures S4A, B,** and **C**). Altogether, these results indicated that Tcb3 is localised in proximity to Sfk1 at the PM, especially in response to heat stress conditions that elevate cytoplasmic Ca^2+^ signalling and induce Tcb3-mediated ER-PM tethering.

**Figure 4.**
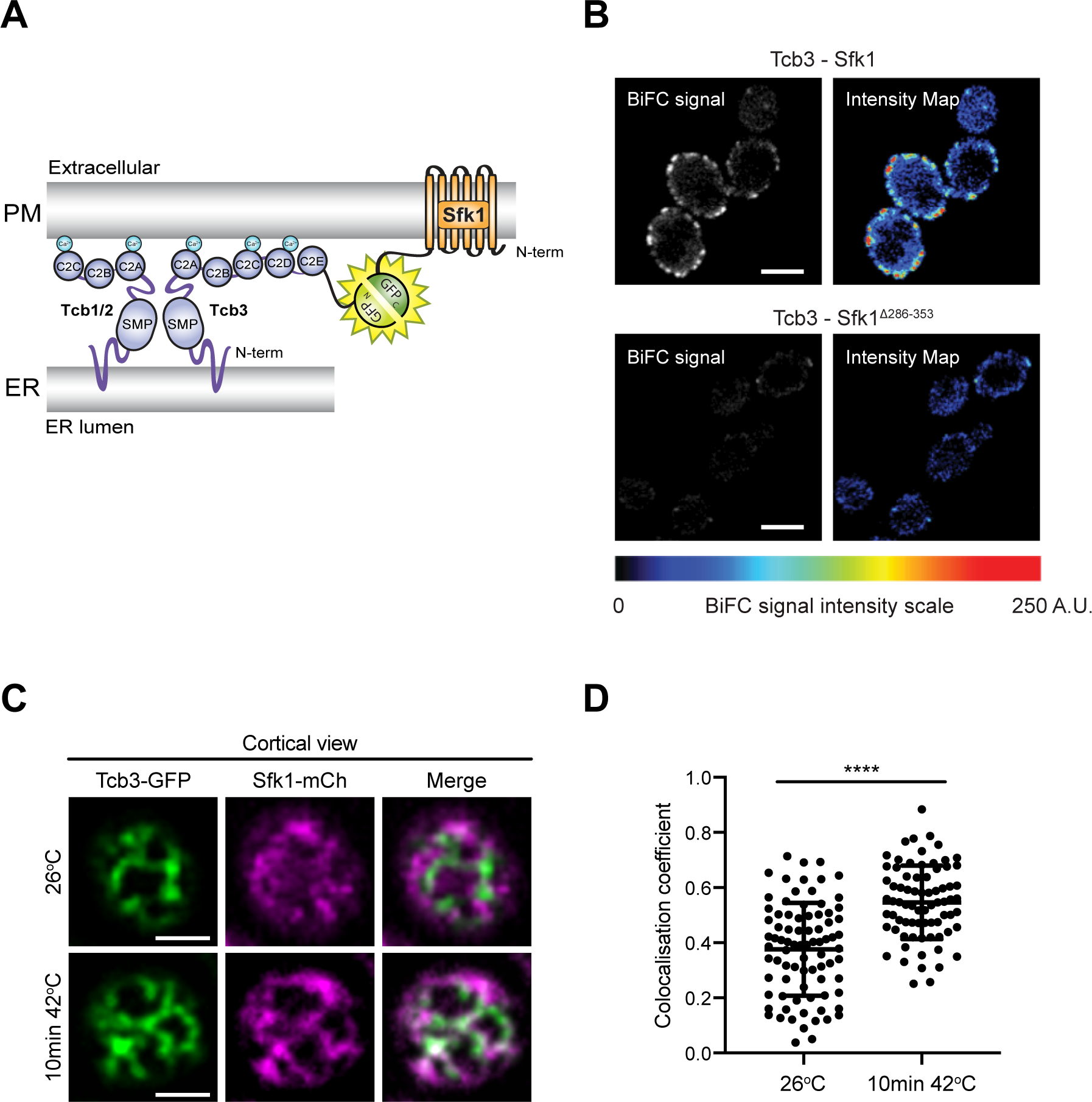
Tcb3 co-localises with the integral PM protein Sfk1. (**A**) Cartoon displaying the bi-molecular fluorescence (BiFC) split GFP assay to assess Tcb3-Sfk1 proximity. The N-terminal half of GFP (GFP_N_) and the C-terminal half of GFP (GFP_C_) form a fluorescent GFP only when their fusion partners, in this case Tcb3 and Sfk1, are in close spatial proximity with each other. (**B**) Tcb3-GFP_N_ associates with Sfk1-GFP_C_ but not with a mutant Sfk1 lacking its cytoplasmic C-terminus (Sfk1^Δ286-353^-GFP_C_). The pseudo-colored images indicate the scale of specific BiFC signals (blue, moderate; red, strong). Scale bars, 4μm (**A**) Cortical localisation of Tcb3-GFP and Sfk1-mCherry at 26°C or after 10 min at 42°C. Scale bars, 4μm (**B**) Quantitation of Tcb3-GFP and Sfk1-mCherry co-localisation (Pearson’s coefficient) at 26°C or after 10 min at 42°C. Data represents mean ± standard deviation. Total number of cells analysed in three independent experiments: wild type n= 82, *tcb1/2/3*Δ n=75). **** p > 0.0001. Also see **Figure S4**.

We next addressed whether the Tcb proteins regulate PS at the PM in response to heat stress. Remarkably, whilst wild type cells showed only a small drop in relative levels of the PS FLARE at the PM after a brief shift from 26°C to 42°C, PM localisation of the PS FLARE significantly decreased in the *tcb1/2/3*Δ mutant cells at 42°C (**Figures 5A** and **B**). A previous study has implicated E-Syt1 in recycling DAG from the PM to the ER during the phosphophoinositide cycle (Saheki et al., 2016). However, in control experiments, the DAG FLARE GFP-C1_PKD_ was not stabilised at the PM in *tcb1/2/3*Δ mutant cells at 26°C or 42°C as compared to wild type control cells (**Figure S5A**). Thus, the Tcb proteins are required for phospholipid, but not glycerolipid, homeostasis at the PM upon heat stress. Regardless, these results do not exclude roles for the Tcb proteins in DAG transport at other inter-organelle contacts or under other stress conditions.

**Figure 5.**
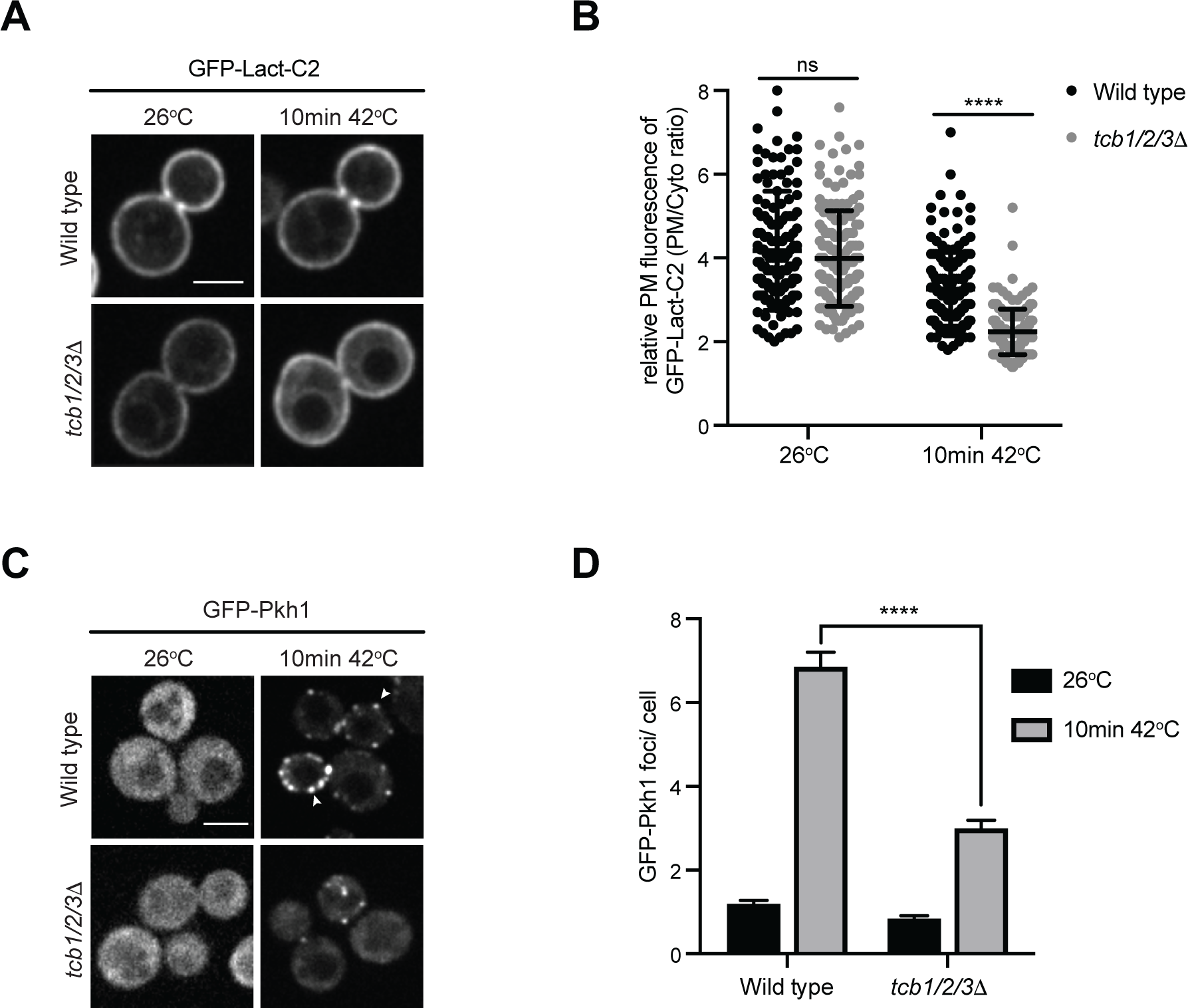
Tricalbin proteins control phosphatidylserine distribution and Pkh1 localisation under heat shock conditions. (**A**) PS FLARE (GFP-Lact-C2) localisation in wild type and *tcb1/2/3*Δ cells at 26°C or after 10 min at 42°C. Scale bar, 4 μm. (**B**) Quantitation of relative GFP-Lact-C2 levels at the PM at 26°C and after 10 min at 42°C in wild type and *tcb1/2/3*Δ cells. Data represents mean ± standard deviation. Total number of cells analysed in three independent experiments: wild type 26°C n=153, wild type 10 min 42°C n=152, *tcb1/2/3*Δ 26°C n=152, *tcb1/2/3*Δ 10 min 42°C n=151 cells. **** p > 0.0001. (**C**) GFP-Pkh1 localisation in wild type and *tcb1/2/3*Δ cells at 26°C or after 10 min at 42°C. White arrows show examples of GFP-Pkh1 cortical patches in wild type cells at 42°C. Scale bar 4μm. (**D**) Quantitation of the number of GFP-Pkh1 foci per cell at 26°C and after 10 min at 42°C in wild type and *tcb1/2/3*Δ cells. Total number of cells analysed in three independent experiments: wild type 26°C n=229, wild type 10 min 42°C n=216, *tcb1/2/3*Δ 26°C n=196, *tcb1/2/3*Δ 10 min 42°C n=173 cells from three independent experiments. **** p > 0.0001. Also see **Figure S5**.

As discussed, the Tcb proteins form heat-induced ER-PM contacts and are required for PM integrity under these conditions (Collado et al., 2019). Yet, how the Tcb proteins contribute to PM integrity is not known. A previous study reported that Pkh1, an ortholog of mammalian phosphoinositide-dependent protein kinase-1 (PDK1), assembles at cortical patches in response to a brief shift to 42°C and is required for PM integrity (Omnus et al., 2016). Pkh1, along with its paralog Pkh2, activates the protein kinases Ypk1 and Ypk2 (AGC kinase family members) that have essential roles in cell signalling, endocytosis, lipid metabolism, and PM integrity (Casamayor et al., 1999; Sun et al., 2000; Friant et al., 2001; Roelants et al., 2002; Roelants et al., 2011; Muir et al., 2014; Omnus et al., 2016). We examined Pkh1 localisation in wild type and *tcb1/2/3*Δ cells using a functional GFP-Pkh1 fusion under non-stress and heat stress conditions (26°C and 42°C). Consistent with a previous study (Omnus et al., 2016), GFP-Pkh1 was diffusely localised in the cytoplasm of wild type cells at 26°C. However, upon a brief heat shock (10 min 42°C), increased numbers of cortical puncta were observed along with some intracellular puncta (**Figures 5C** and **D**). The number of cortical GFP-Pkh1 foci per cell after heat shock was significantly lower in *tcb1/2/3*Δ mutant cells (>2-fold; **Figures 5C** and **D**). Surprisingly, a previous study found that Δtether cells displayed constitutive recruitment of Pkh1 even under normal growth conditions (Omnus et al., 2016). However, this apparent discrepancy may be attributed to increased levels of PI4P in the PM of the Δtether cells but not *tcb1/2/3*Δ cells (see **Figure 2**) that may be detected by the PH-like domain of Pkh1 (**Figure S5B**) (Fidler et al., 2016). The precise lipid-binding specificity of the Pkh1 PH-like domain will require further examination in future studies. However, consistent with a role of phospholipids in Pkh1 regulation, heat-induced Pkh1 puncta formation was significantly impaired in cells lacking Sfk1 (**Figure S5C**). As discussed, Sfk1 has been previously implicated in the control of lipid flip and/or flop in the PM bilayer (Mioka et al., 2018). Accordingly, cells lacking PM phospholipid flippases (*dnf1/2/3*Δ) or floppases (*pdr5*Δ *snq2*Δ *yor1*Δ) displayed opposite effects on GFP-Pkh1 cortical assemblies at 42°C (significantly decreased and increased, respectively; **Figure S5D**).

Next, we compared the relative roles of the ER-PM tethering proteins in PM homeostasis upon heat-induced membrane stress. Notably, PM levels of the PS FLARE in *tcb1/2/3*Δ cells at 42°C resembled those of the Δtether cells at 26°C and 42°C (**Figure 6A**). In other words, loss of Scs2/22 and Ist2 was not additive with loss of Tcb1/2/3 at 42°C. This finding further suggested that Tcb proteins serve a primary role in PS homeostasis at the PM upon heat stress conditions, while Scs2/22 and Ist2 function under non-stress conditions (**Figure 3**). A previous study found that Tcb3 was required for the formation of heat-induced ER-PM contacts (Collado et al., 2019). We therefore examined roles of the Tcb3 protein and its domains in PM phospholipid homeostasis and integrity upon heat-induced membrane stress. Loss of Tcb3 alone resulted in a measurable decrease in PM levels of the PS FLARE, as compared to wild type cells at 42°C (1.5-fold; **Figure 6B**). This was rescued by expression of a Tcb3-GFP fusion from a plasmid, but not by a mutant form of Tcb3 lacking the SMP domain (Tcb3ΔSMP-GFP) (**Figure 6B**) or an artificial tether (GFP-MSP-Sac1) (**Figure 6B**) previously shown to restore ER-PM contacts in the Δtether cells (Manford et al., 2012). Accordingly, loss of Tcb3 alone resulted in significant PM integrity defects at 42°C, as measured by an established quantitative assay (>10-fold increase in propidium-stained *tcb3*Δ cells; **Figure 6C**), consistent with a previous study (Collado et al., 2019). This was efficiently rescued by expression of Tcb3-GFP, but not by mutant forms of Tcb3 lacking either the SMP or C2 domains (Tcb3ΔSMP-GFP or Tcb3ΔC2-GFP, respectively) (**Figure 6C**) or by the artificial tether (GFP-MSP-Sac1) (**Figure 6C**). Finally, cells lacking the Tcb proteins displayed significant increases in the amplitude and duration of cytoplasmic Ca^2+^ bursts at 42°C, as compared to wild type cells (**Figures S6A, B,** and **C**). Altogether, our findings indicate that the Tcb proteins are Ca^2+^-activated lipid transfer proteins that maintain PM integrity and modulate cytoplasmic Ca^2+^ signalling upon stress conditions.

**Figure 6.**
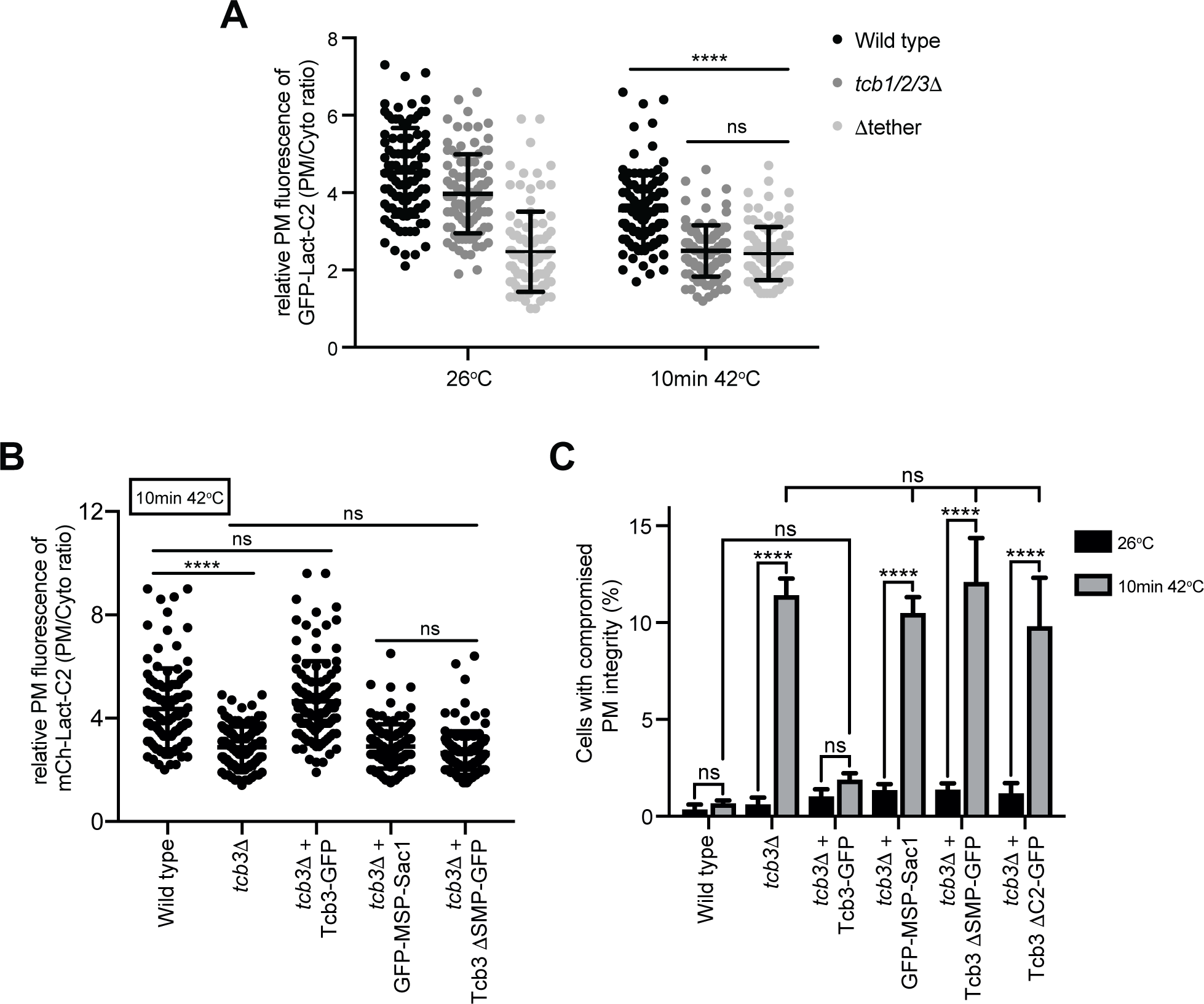
The SMP domain of Tcb3 is required for phosphatidylserine homeostasis and PM integrity under heat shock conditions. (**A**) Quantitation of relative GFP-Lact-C2 levels at the PM in wild type, *tcb1/2/3*Δ, and Δtether cells at 26°C or after 10 min at 42°C. Data represents mean ± standard deviation. Total number of cells analysed in three independent experiments: all strains and conditions n=100. **** p > 0.0001; ns, not significant. (**B**) Quantitation of relative GFP-Lact-C2 levels at the PM in wild type, *tcb3*Δ, and *tcb3*Δ cells expressing Tcb3-GFP, a mutant Tcb3 fusion lacking the SMP domain (Tcb3ΔSMP-GFP), or an artificial ER-PM tether (GFP-MSP_Scs2_-Sac1) (Manford et al., 2012) after 10 min at 42°C. Data represents mean ± standard deviation. Total number of cells analysed in three independent experiments: wild type n=108, *tcb3*Δ n=116, *tcb3*Δ + Tcb3-GFP n=108, *tcb3*Δ + Tcb3ΔSMP-GFP n=112, *tcb3*Δ + GFP-MSP_Scs2_-Sac1 n=112). **** p > 0.0001; ns, not significant. (**C**) PM integrity assays of wild type, *tcb3*Δ and *tcb3*Δ cells complemented with Tcb3-GFP, Tcb3-GFP truncation mutants (Tcb3ΔSMP-GFP or Tcb3ΔC2-GFP), or GFP-MSP_Scs2_-Sac1. Cells incubated at 26°C or 42°C for 10 min were subsequently incubated with propidium iodide and measured by flow cytometry (50,000 cells measured per experiment). Data represents mean ± standard deviation from three independent experiments. **** p > 0.0001; ns, not significant. Also see **Figure S6**.

## Discussion

The primary functions of the evolutionarily conserved Tcb/E-Syt family members have remained elusive. This has been partly due to the lack of severe phenotypes observed upon loss of the E-Syt proteins. For example, previous studies reported no obvious defects in triple E-Syt1/2/3 knockout mice (Sclip et al., 2016; Tremblay and Moss, 2016). However, the E-Syt/Tcb proteins have been conserved across species throughout evolution (Lee and Hong, 2006; Kopec et al., 2010; Wong and Levine, 2017), suggesting they must serve important functions that have been overlooked. Indeed, studies in yeast have revealed roles for the Tcb proteins in PM integrity under membrane stress conditions (Aguilar et al., 2007; Omnus et al., 2016; Collado et al., 2019) (see **Figures 6** and **7**). Likewise, Esyt2/Esyt3 double knockout mouse embryonic fibroblasts displayed decreased migration and survival under *in vitro* stress culture conditions (Herdman et al., 2014). Furthermore, E-Syt family members have been implicated in neuronal growth and survival (Kikuma et al., 2017; Gallo et al., 2020; Nath et al., 2020). However, mechanistic detail on E-Syt function in these essential processes has been lacking. Using quantitative sensors and lipidomics, we have found that the Tcb proteins control phospholipid homeostasis at the PM. Phospholipid regulation may be an anciently conserved role of the E-Syt protein family in maintaining cellular homeostasis.

**Figure 7.**
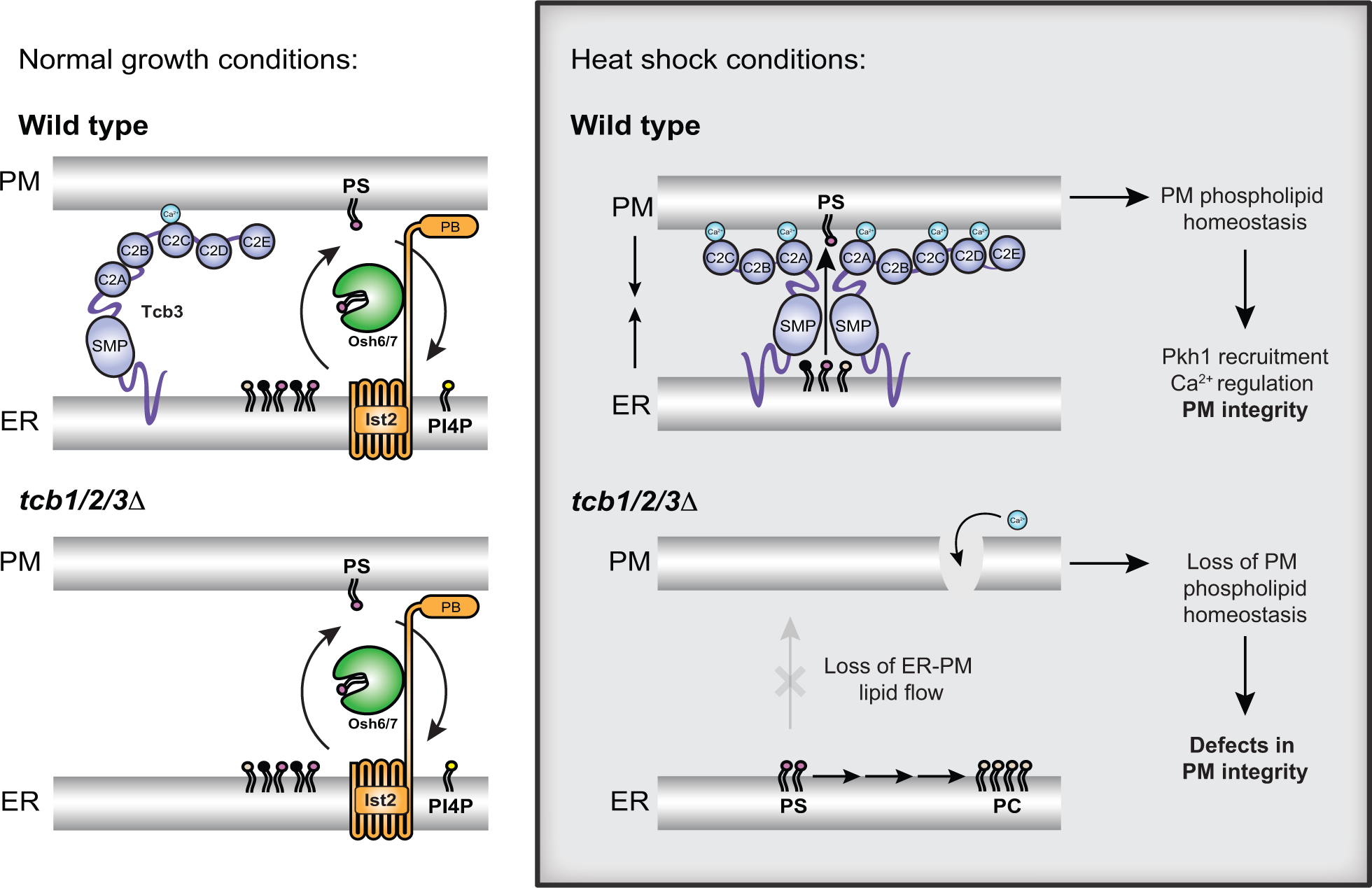
Speculative model for tricalbin protein function in PM phospholipid homeostasis and integrity. Under normal growth conditions (left panels), non-vesicular phosphatidylserine (PS, magenta) transport from the ER to the PM is carried out by Osh6 and Osh7 that are recruited to ER-PM contacts by Ist2 (D’Ambrosio et al., 2020). Osh6 and Osh7 move PS from the ER to PM in exchange for PI4P (yellow) at the PM (top left, Wild type cells). Under normal growth conditions, the tricalbin (Tcb) proteins may have a structural role in tethering the ER and PM, but they are not required for PS transport as loss of Tcb1/2/3 does not substantially affect PS distribution (bottom left, *tcb1/2/3*Δ cells). Under heat shock conditions, however, increases in cytoplasmic Ca^2+^ levels further induce Tcb3 function. This may include dimerisation with Tcb1/2 and interactions with the PM through the C2 domains that may promote the ability of the Tcbs to form heat-induced ER-PM contacts (Collado et al., 2019). Importantly, under heat shock conditions, the Tcbs serve a primary role in PS homeostasis at the PM, either by SMP domain-mediated non-vesicular PS transfer from the ER to the PM (top right) or by recruiting additional PS transfer proteins to ER-PM contacts (such as Osh6/7, not depicted). Loss of the Tcb proteins (bottom right, *tcb1/2/3*Δ) leads to a lack of PS delivery to the PM in heat stressed cells, resulting in loss of Pkh1 recruitment (see Figure 5), as well as prolonged Ca^2+^ bursts and PM integrity defects (bottom right, *tcb1/2/3*Δ).

Our results indicate that the Tcb proteins control levels of phospholipids that are enriched at the PM. In particular, localisation of a PS-specific reporter was reduced at the PM upon loss of the Tcb proteins (**Figures 3B, 3C, 5A, 5B, 6A,** and **6B**). PS has essential roles in PM organisation in eukaryotic cells (Yeung et al., 2008; Fairn et al., 2011; Cho et al., 2016; Haupt and Minc, 2017; Sartorel et al., 2018). It is intriguing that previous studies had not uncovered clear roles for the E-Syt proteins in PS regulation. However, one study found that whilst mice lacking all three E-Syt were viable, the PS transfer proteins ORP5 and ORP8 were upregulated (Tremblay and Moss, 2016), providing a potential compensatory mechanism. Another study found no differences in PS levels in isolated PM fractions between E-Syt1/2/3 triple knockout and control cells by lipidomics (Saheki et al., 2016); however, this experiment was performed at basal cytoplasmic Ca^2+^ concentrations where E-Syt activity is low. More recently, the E-Syt proteins have been implicated in Ca^2+^-mediated PS externalisation (Bian et al., 2018), and this could be attributed to impaired PS delivery to the PM. Yet, potential roles of the E-Syt proteins in PS regulation in mammalian cell remain to be explored.

Our quantitative lipidomics results provide direct evidence that the Tcb proteins control phospholipid homeostasis at the PM. Previous studies have reported alterations in phospholipid levels in Δtether yeast cells lacking several ER-PM tether proteins (Quon et al., 2018; Nishimura et al., 2019; Jorgensen et al., 2020; Wang et al., 2020). However, none of these studies addressed the specific contribution of the Tcb proteins. Two studies even questioned the roles of ER-PM contacts in lipid transfer (Quon et al., 2018; Wang et al., 2020). However, neither of these studies analysed acyl chain composition that provides important information on lipid localisation (*e.g.* ER *versus* PM species). Our species-level lipidomic results show that mono-unsaturated PS and PE, most notably 32:1 and 34:1 isoforms shown to be enriched at the PM (Schneiter et al., 1999), are specifically depleted upon loss of the Tcb proteins (**Figures 3D, 3E,** and **S3B**). However, there was no decrease in di-unsaturated forms of PS and PE (**Figure S3B**). The specific loss of PM phospholipid species is consistent with a role of the Tcb proteins in coordinating phospholipid synthesis and transfer at ER-PM contacts. All E-Syt family members feature a SMP domain that dimerises and transports glycerolipids *in vitro* (Lee and Hong, 2006; Toulmay and Prinz, 2012; Schauder et al., 2014; Saheki et al., 2016; Yu et al., 2016; Bian et al., 2018; Bian and De Camilli, 2019; Qian et al., 2021). In particular, a recent study demonstrated the ability of the Tcb3 SMP domain to transport phospholipids *in vitro* (Qian et al., 2021), consistent with our *in vivo* experiments. Furthermore, the SMP domain is required for Tcb protein localisation and PM integrity (Toulmay and Prinz, 2012; Collado et al., 2019) (**Figure 6C**), suggesting that the SMP domain-containing Tcb proteins may directly deliver mono-unsaturated phospholipids to the PM at ER-PM contacts **(Figure 7)**. Although, the data do not exclude the possibility that Tcb proteins may serve as tethers for additional lipid transfer proteins at ER-PM contacts.

E-Syt family members have been implicated in distinct steps of the phosphoinositide cycle in metazoan cells. For example, E-Syt proteins have been suggested to facilitate the activity of certain phosphatidylinositol transfer proteins (PITPs) at ER-PM contacts, including Nir2/3 and Rgdβ in mammalian cells and *Drosophila* respectively (Chang et al., 2013; Nath et al., 2020). The Nir2/3 and Rgdβ PITPs are proposed to transfer PI from the ER to the PM for the generation of PI4P and PI(4,5)P_2_ at the PM. However, PI4P synthesis at the PM is not impaired upon loss of the Tcb proteins or other ER-PM tethers, Scs2/22 and Ist2 (**Figures 2B, 2C, 2E** and **S2B**) (Manford et al., 2012). Thus, the Tcb proteins and other ER-PM contact proteins (Scs2/22 and Ist2) are not required for delivery of PI to the PM in yeast. Furthermore, orthologs of the Nir2/Rgdβ PITPs do not exist in fungi (Hsuan and Cockcroft, 2001), and so putative roles of E-Syt proteins in PI transfer may have evolved separately in animals. Mammalian cells lacking the E-Syt proteins also feature prolonged accumulation of DAG at the PM following phospholipase C (PLC)-mediated PI(4,5)P_2_ hydrolysis (Saheki et al., 2016), suggesting a defect in recycling DAG to the ER during the phosphoinositide cycle. However, we did not observe accumulation of DAG at the PM in cells lacking the Tcb proteins, using a *bona fide* sensor of Plc1-generated DAG (**Figure S5A**). This result does not rule out a potential role for the Tcbs, or other SMP domain-containing proteins, in DAG channelling at intra-organelle contacts in yeast. However, given the apparent lack of effects on PI and DAG and the major changes in PS and PE, the Tcb proteins may primarily regulate PS and PE homeostasis at the PM.

While the Tcb proteins and ER-PM contacts are not required for PI transfer, our study does reveal links to PI kinase activity at the PM. In particular, ER-localised Tcb3 is found in proximity to the PM-localised Sfk1 protein (**Figure 4**). Sfk1 (TMEM150 ortholog) is purportedly a subunit of a PI 4-kinase (PI4K) complex that generates PI4P at the PM and is required for heat-stimulated PI(4,5)P_2_ synthesis (Audhya and Emr, 2002). Moreover, Tcb3 and Sfk1 co-localisation increases following heat shock (**Figure 4B**). Steady-state lipidomics measurements reveal that while PI4P is elevated upon loss of the Tcb proteins, PI(4,5)P_2_ levels are not (**Figure 2B, 2D, 2F** and **S2B**), suggesting that the Tcbs may contribute to the conversion of PI4P to PI(4,5)P_2_. PIP 5-kinase is recruited and activated by PS (Fairn et al., 2009; Nishimura et al., 2019), and regulation of PS by the Tcb proteins may be involved. Interestingly, Sfk1 is also implicated in PM phospholipid asymmetry and PM integrity (Mioka et al., 2018). As PI(4,5)P_2_ synthesis is required for PM integrity (Audhya et al., 2000; Omnus et al., 2016), one possibility is that Tcb3 and Sfk1 promote the formation of PS-enriched domains that stimulate PIP5K activity upon PM stress. In support of this, the Tcb proteins are required for PS homeostasis at the PM upon heat shock (**Figures 5A, 5B, 6A,** and **6B**). Accordingly, the Tcb proteins are activated upon heat-induced Ca^2+^ signalling and form distinct ER-PM contacts that are required for PM integrity under these conditions (**Figures S4** and **6C**) (Omnus et al., 2016; Collado et al., 2019).

We also show that the Tcb proteins facilitate heat-induced PM recruitment of the protein kinase Pkh1, a PDK1 ortholog necessary for PM integrity upon heat shock (**Figures 5C** and **D**) (Roelants et al., 2002; Roelants et al., 2011; Omnus et al., 2016). Both PI4P and PI(4,5)P_2_ are implicated in activation of Pkh1 at the PM upon stress conditions (Omnus et al., 2016). The carboxy-terminal region of Pkh1 contains a PH domain-like fold (Fidler et al., 2016) and PH domains often serve as co-incidence detectors of phosphoinositides and PS (Vonkova et al., 2015; Nishimura et al., 2019). Accordingly, mammalian PDK1 is regulated by both PI(3,4,5)P_3_ and PS (Lucas and Cho, 2011; Heras-Martínez et al., 2019). Here, we find that *tcb1/2/3*Δ cells have reduced levels of a PS FLARE at the PM after heat shock (**Figures 5A** and **B**). Correct maintenance of phospholipids within the PM may be essential for Pkh1 PM recruitment. Consistent with this, loss of PM lipid flippases and floppases modulated Phk1 cortical assembly upon heat shock (**Figure S5**). Accordingly, Tcb-mediated PS regulation may be critical for Pkh1 localisation and PM integrity. However, our results do not rule out roles of other lipids in PM maintenance and integrity. For example, the Tcb proteins have recently been implicated in non-vesicular transport of ceramide from the ER to Golgi compartments (Ikeda et al., 2020).

Correct maintenance of PS within the PM is integral to cell survival (Matsuo et al., 2007) and in mammalian cells, rearrangements in PS distribution within the PM bilayer acts as a trigger for apoptotic cell death (Fadok et al., 1992). PS also accumulates at sites of PM damage (Horn and Jaiswal, 2019) and is involved in the recruitment of Ca^2+^- and PS-binding proteins such as annexins, dysferlins, and synaptotagmins during PM repair (Middel et al., 2016; Boye et al., 2017; Horn and Jaiswal, 2018). However, the mechanism for PS enrichment at sites of PM damage is incompletely understood. Regulation of PS dynamics may be an anciently conserved function of E-Syt family members in PM integrity and repair. The synaptotagmin proteins have been the subject of intense focus due to their functions in neurotransmitter release at neuronal synapses (Südhof, 2002; Chapman, 2008), although they also act as Ca^2+^-dependent regulators of exocytic events in both neuronal and non-neuronal exocrine cells, driving SNARE-mediated membrane fusion (Hui et al., 2009; van den Bogaart et al., 2011; Kim et al., 2012). Accordingly, synaptotagmins and dysferlins are proposed to aid in PM repair by triggering vesicle fusion at sites of injury (Detrait et al., 2000; Lek et al., 2013). Synaptotagmin family members are not conserved in fungi, unlike the extended-synaptotagmins (E-Syts) that are expressed ubiquitously throughout the eukaryotic as well as prokaryotic kingdoms (Wong and Levine, 2017). This hints to a more ancient fundamental function of the E-Syt proteins that predates that of the C2 domain-containing synaptotagmins. Protein kinase C isoforms that bear a C2 domain may be a common ancestral effector protein involved in cellular integrity and exocytosis in yeast and mammalian cells (Levin, 2005; Horn and Jaiswal, 2018; Horn and Jaiswal, 2019). Therefore, the E-Syt family may confer a primordial mechanism for PM maintenance that subsequently evolved in higher eukaryotes in the control of regulated exocytosis during PM wound repair and eventually even insulin and neurotransmitter release.

The importance of Ca^2+^ signalling in membrane stress responses cannot be understated. Any weakening or disruption of the PM leads to influx of the ion into the cytoplasm, resulting in an increase in cytoplasmic Ca^2+^ concentration up to two orders of magnitude (from 0.1 μM to up to 10 μM) (Jaiswal, 2001). Whilst Ca^2+^ is recognised as a signal for membrane damage response and repair pathways, increases in cytoplasmic Ca^2+^ above a defined threshold (> 10 μM) can result in loss of membrane organelle and cellular integrity leading to cell death, as originally described by Zimmerman and Hülsmann as “the calcium paradox” (Zimmerman and Hülsmann, 1966). Accordingly, cells have established rapid response systems to damage- and stress-induced Ca^2+^ influx. Many of the proteins involved in PM repair mechanisms (including exocytosis-mediated vesicle fusion, endocytosis, membrane shedding, and reorganisation of the cytoskeleton in response to PM stress or injury) are Ca^2+^-binding proteins that function as Ca^2+^ sensors (Benink and Bement, 2005; Cheng et al., 2014; Holmes et al., 2015; Horn and Jaiswal, 2018; Koerdt et al., 2019). The E-Syt family members are no exception and function as sensors of large increases in cytoplasmic Ca^2+^ (> 1 μM) (Idevall-Hagren et al., 2015). Accordingly, mammalian E-Syt proteins are regulated by the store-operated Ca^2+^ entry (SOCE) pathway and subsequently inactivate SOCE (Giordano et al., 2013; Idevall-Hagren et al., 2015). We find that the yeast Tcbs are active under conditions that increase cytoplasmic Ca^2+^ signals (**Figures 1** and **S4**) and the Tcbs in turn attenuate cytoplasmic Ca^2+^ signalling (**Figure S6**). Thus Tcb/E-Syt function at ER-PM contacts responds to and modulates cytoplasmic Ca^2+^ signals, and this regulatory system is conserved from yeast to humans.

In summary, we propose that the yeast Tcb proteins serve as Ca^2+^-activated ER-PM tethers and lipid transfer proteins that maintain PS levels necessary for PM integrity under stress conditions (**Figure 7**). Tcb-mediated regulation of PS in the cytosolic leaflet of the PM aids in the direct recruitment of protein and lipid kinases required for PM integrity. Essential roles of the mammalian E-Syt proteins in cell and tissue homeostasis may be revealed under conditions of PM stress conditions, and it will be interesting to re-examine the E-Syt proteins in this context.

## Materials and Methods

### Yeast Strains, Plasmids, Media and Growth Assays

Descriptions of strains and plasmids used in this study used in this study are listed in **Supplemental Tables 1** and **2**. Gene deletions and epitope tags were introduced into yeast by homologous recombination (Longtine et al., 1998). The pRS vector series have been described previously (Sikorski and Hieter, 1989). Plasmids were sequenced to ensure that no mutations were introduced due to manipulations. Standard techniques and media were used for yeast and bacterial growth. For plating assays in **Figure S1**, cells were grown to midlog, adjusted to 1 OD600/ml, and serial dilutions were plated on agar media either containing or lacking 0.75 μM myriocin. Haploid Split-GFP strains listed in **Supplemental Table 1** were made using tagging cassettes (Barnard et al., 2008). Diploid strains were then created by mating to obtain various combinations of N- and C-terminal GFP fragment-tagged target proteins.

### Live Yeast Cell Imaging

Fluorescence microscopy experiments were performed on mid-log yeast cultures in synthetic media at the indicated temperatures. Live yeast cell imaging data in all Figures were acquired with a 100× CFI Plan Apochromat VC oil-immersion objective lens (1.4 NA), using a PerkinElmer Ultraview Vox spinning disk confocal microscope that consists of a Nikon TiE inverted stand attached to a Yokogawa CSU-X1 spinning disk scan head, a Hamamatsu C9100-13 EMCCD camera, Prior NanoscanZ piezo focus, and a Nikon Perfect Focus System (PFS). All images were collected as square images with 512 x 512 pixels. The number of cells observed in experiments are reported in the figures and figure legends. The brightness and contrast of images were linearly adjusted and cropped in Photoshop (Adobe) for presentation.

### Quantitative Image Analysis

All quantitative image analyses were conducted using ImageJ/Fiji (Schindelin et al., 2012).

To calculate the specific PM to cytosol (PM/Cyto) ratio of specific lipid species in **Figures S1, 2, 3, 5** and **6**, the relative fluorescence (relative F_PM_) was quantified as described in (Nishimura et al., 2019). Briefly, individual cells were chosen from single channel images, lines were drawn cross the mother cell and the corresponding fluorescence intensity profiles were plotted. The two highest intensity values, corresponding to signal at the PM, were averaged (F_PM_). Intensity measurements were also taken from lines drawn through the cytosol (F_cytosol_) and background (F_background_) and PM relative fluorescence was calculated by using the equation: relative F_PM_ = (F_PM_-F_background_)/(F_cytosol_-F_background_). Peaks in intensity profiles were automatically calculated by an Excel VBA macro.

Split GFP (Barnard et al., 2008) signal intensity in **Figures 4** and **S4** was measured using Fiji. To identify GFP-Pkh1 puncta per cell in **Figures 5** and **S5**, individual cells were selected and for each cell, points of interest (GFP-Pkh1 foci) were identified using the Find Maxima tool in Fiji applying appropriate noise tolerance settings.

### LC-MS/MS analysis of methylated PIPs

Levels of individual PI, PIP and PIP2 species in wild type, *tcb1/2/3*Δ, *scs2/22*Δ *ist2*Δ and Δtether mutant cell extracts **Figures 2** and **S2** were analysed as previously described (Nishimura et al., 2019). 20 OD_600_ units of cells were precipitated and washed with cold 4.5% perchloric acid. For phosphoinositide measurements, cells were resuspended in 500 µL 0.5 M HCl and disrupted with a 5.0 mm zirconia bead by vigorous shaking (1,500 rpm for 10 min) using Shake Master Neo (BMS, Tokyo, Japan). The homogenates were transferred to new tubes and centrifuged at 15,000 × *g* for 5 min. The pellets were resuspended in 170 μL water and 750 μL of CHCl_3_/MeOH/1 M HCl (2:1:0.1, v/v) and incubated for 5 min at room temperature. To each sample 725 μl of CHCl_3_ and 170 μl of 2 M HCl were added, followed by vortexing. After centrifugation at 1,500 × *g* for 5 min, the lower phase was collected and washed with 780 μl of pre-derivatization wash solution (the upper phase of CHCl_3_/MeOH/0.01 M HCl (2:1:0.75 v/v)). The lipid extracts were derivatized by adding 50 μl of 2 M TMS-diazomethane in hexane. The derivatization was carried out at room temperature for 10 min and was stopped by adding 6 μl of glacial acetic acid. The derivatized samples were washed twice with 700 μl of post-derivatization wash solution (the upper phase of CHCl_3_/MeOH/water (2:1:0.75 v/v)). After adding 100 μl of MeOH/H_2_O (9:1, v/v), the samples were dried under a stream of N_2_, dissolved in 80 μl of MeOH and sonicated briefly. After adding 20 μl of water, the samples were subjected to LC-ESI-MS/MS analysis. The LC-ESI-MS/MS analysis was performed on a Shimadzu Nexera ultra high performance liquid chromatography system coupled with a QTRAP 4500 hybrid triple quadrupole linear ion trap mass spectrometer. Chromatographic separation was performed on an Acquity UPLC C4 BEH column (100 mm × 1 mm, 1.7 μm; Waters) maintained at 40°C using mobile phase A (water containing 0.1% formate) and mobile phase B (acetonitrile containing 0.1% formate) in a gradient program (0–5 min: 45% B; 5–10 min: 45% B→100% B; 10-15 min: 100% B; 15–16 min: 100% B→45% B; 16-20: 45% B) with a flow rate of 0.1 mL/min. The instrument parameters for positive ion mode were as follows: curtain gas, 10 psi; collision gas, 7 arb. unit; ionspray voltage, 4500 V; temperature, 600°C; ion source gas 1, 30 psi; ion source gas 2, 50 psi; declustering potential, 121 V; entrance potential, 10 V; collision energy, 39 V; collision cell exit potential, 10 V. Methylated phosphoinositides and phosphatidylserine were identified and quantified by multiple reaction monitoring. For these measurements, internal standards of 10 ng of 17:0-20:4 PI, PI(4)P, and PI(4,5)P_2_, and PS were added to each sample.

### Quantitative shotgun lipid MS data acquisition, analysis, and post-processing

Levels of individual PS, PE and PC species in wild type, *tcb1/2/3*Δ, *scs2/22*Δ *ist2*Δ and Δtether mutant cell extracts in **Figures 3** and **S3** were determined by mass spectrometry-based quantitative, shotgun lipidomics by Lipotype GmbH (Dresden, Germany) as described (Ejsing et al., 2009; Klose et al., 2012). Total yeast cell lysate samples were diluted to 0.2 OD units using 155 mM ammonium bicarbonate in water to the total volume of 150 µl and were spiked with internal lipid standard mixture. Lipids were extracted using a two-step chloroform/methanol procedure with 750 µl volume of each organic phase step (chloroform:methanol, 15:1 and 2:1 respectively for the 1^st^ and the 2^nd^ step)(Ejsing et al., 2009). After extraction, the organic phase was transferred to an infusion plate and dried in a speed vacuum concentrator. 1^st^ step dry extract was re-suspended in 100 µl 7.5 mM ammonium acetate in chloroform/methanol/propanol (1:2:4, V:V:V) and 2^nd^ step dry extract in 100 µl 33% ethanol solution of methylamine in chloroform/methanol (0.003:5:1; V:V:V). All liquid handling steps were performed using Hamilton Robotics STARlet robotic platform with the Anti Droplet Control feature for organic solvents pipetting. Samples were analysed by direct infusion on a QExactive mass spectrometer (Thermo Scientific) equipped with a TriVersa NanoMate ion source (Advion Biosciences). Samples were analysed in both positive and negative ion modes with a resolution of R_m/z=200_=280000 for MS and R_m/z=200_=17500 for MSMS experiments, in a single acquisition. MSMS was triggered by an inclusion list encompassing corresponding MS mass ranges scanned in 1 Da increments (Surma et al., 2015). Both MS and MSMS data were combined to monitor EE, DAG and TAG ions as ammonium adducts; PC as an acetate adduct; and PA, PE, PG, PI and PS as deprotonated anions. Data were analysed with in-house developed lipid identification software based on LipidXplorer (Herzog et al., 2011; Herzog et al., 2012). Data post-processing and normalization were performed using an in-house developed data management system. Only lipid identifications with a signal-to-noise ratio >5, and a signal intensity 5-fold higher than in corresponding blank samples were considered for further data analysis

### Quantitative GCaMP3 Fluorescence Assays

To quantify resting cytosolic Ca^2+^ levels in **Figure 1**, strains expressing a cytoplasmic GCaMP3 reporter were grown at 26°C. For flow cytometry experiments, cells were transferred to PBS. Mean fluorescence of 50,000 events was recorded on a BD Accuri C6 flow cytometer. Background was determined using strains harbouring vector alone. For fluorescence microscopy measurements cells were transferred immediately to a slide and imaged directly. For identification of Ca^2+^ bursts after heat shock **in Figures S4** and **S6**, cells were incubated for 10 min at the indicated temperature before being directly transferred to a slide and imaged at max speed for 5 minutes. Individual cells were chosen from single channel time lapse images. Intensity measurements were taken in an area of the same size within the cell (F_cell_) and adjacent background (F_background_). The normalised GCaMP3 signal (ΔF/F_0_) was calculated from time lapse images, using the equation: ΔF/F_0_ = (F_cell_ at time *t* - F_background_ at time *t*)/ (F_cell_ at time *t=0* - F_background_ at time *t=0*). Peaks in GCaMP3 fluorescence intensity, representing Ca^2+^ bursts, were identified using Prism 8 Area under curve function.

### Plasma membrane Integrity Assay

PM integrity assays in **Figure 6** were performed as described (Omnus et al., 2016; Collado et al., 2019). Yeast strains were grown at 26°C to midlog phase, then kept at 26°C or shifted to either 42°C for 15 min. 1 OD_600_ equivalent of cells was pelleted, resuspended in PBST (0.01% Tween 20), and stained with propidium iodide (Sigma) for 15 min. Cells were then washed twice with PBST and analysed by flow cytometry on a BD Accuri C6 flow cytometer. For flow cytometry analysis, 50,000 cells were counted for each sample from three independent experiments and combined for a total analysis of 150,000 cells. Background was determined by analysing each of the cell strains at the indicated temperatures prior to staining with propidium iodide.

### Quantification and Statistical Analysis

Statistical analysis was carried out using GraphPad Prism 8. To compare the mean of two groups, an unpaired two-tailed t test was used. To compare the mean of multiple groups, we used one-way ANOVA followed by Tukey-Kramer multiple comparisons.

## Acknowledgements

We thank Scott Emr, Jeremy Thorner, Todd Graham, Yuxin Mao, and Greg Fairn for strains and plasmids. We also thank Ruben Fernandez-Busnadiego, Robbie Loewith, Tim Levine, Taki Nishimura, and Bailey Hewlett for helpful discussions. C.J.S. is supported by MRC funding to the MRC LMCB University Unit at UCL, award code MC_UU_00012/6.

## Availability of data and materials

Materials used in this study are available from the corresponding author on request.

## Contributions

F.B.T., D.J.O., J.M.B., G.H.C.C., N.K., and C.J.S. designed and performed experiments; F.B.T., N.K., and C.J.S. analysed the data and wrote the manuscript. All authors discussed the results and commented on the manuscript.

## Additional Data Files

### Supplemental Figure Legends

**Figure S1.**
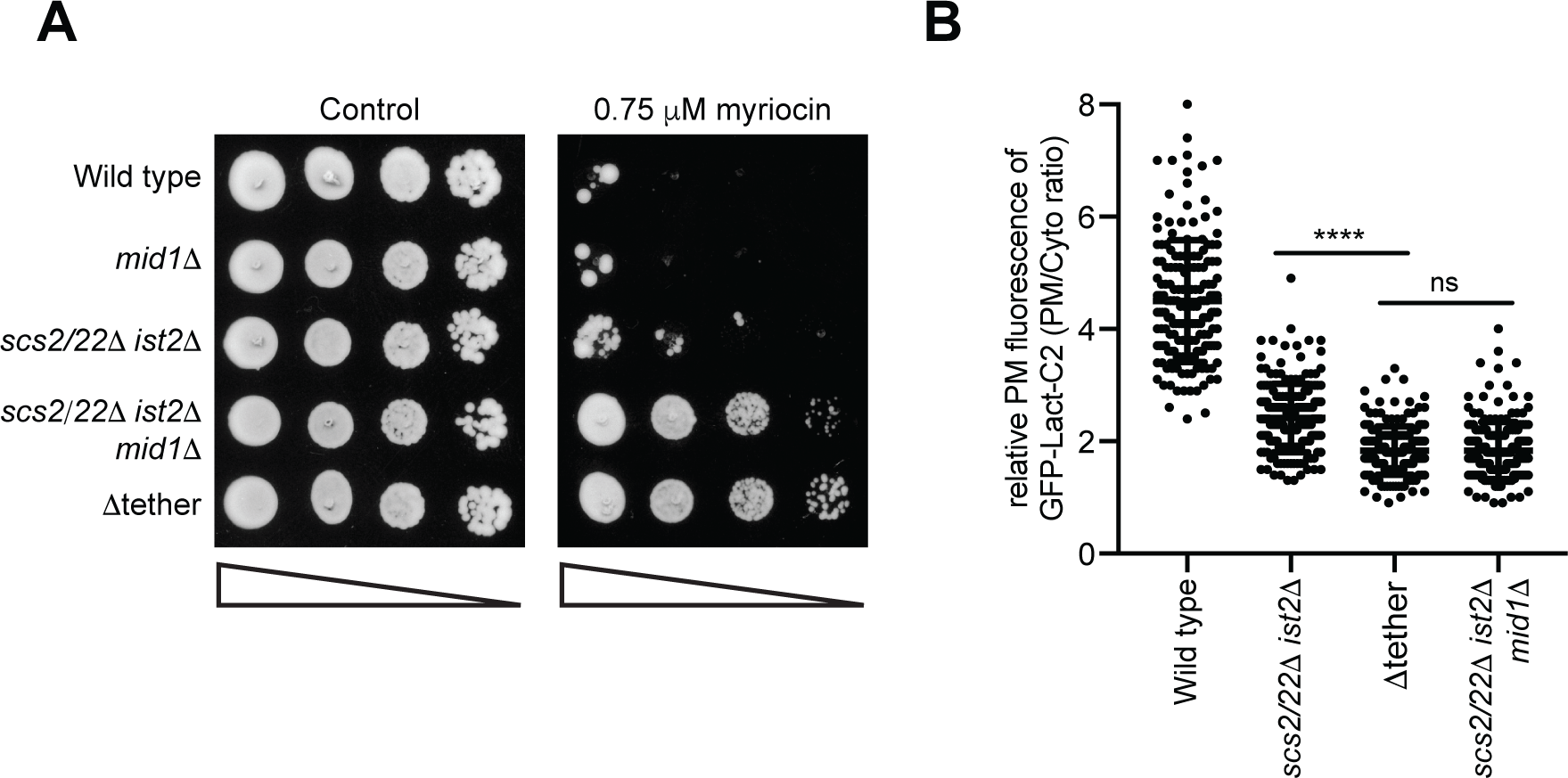
Deletion of the stretch-activated Ca^2+^ channel Mid1 in *scs2/22*Δ *ist2*Δ cells phenocopies the Δtether mutant cells. (**A**) Serial dilutions (10-fold) of wild type, *mid1*Δ, *scs2/22*Δ *ist2*Δ, *scs2/22*Δ *ist2*Δ *mid1*Δ and Δtether cells spotted on agar media +/− 0.75μM myriocin. (**B**) Quantitation of the PS FLARE (GFP-Lact-C2) intensity at the PM of wild type, *scs2/22*Δ *ist2*Δ, Δtether, and *scs2/22*Δ *ist2 mid1*Δ cells. Data represents mean ± standard deviation. Total number of cells analysed in three independent experiments: wild type n=172, *scs2/22*Δ *ist2*Δ n=178, Δtether n=177, *scs2/22*Δ *ist2*Δ *mid1*Δ n=181. **** p > 0.0001.

**Figure S2.**
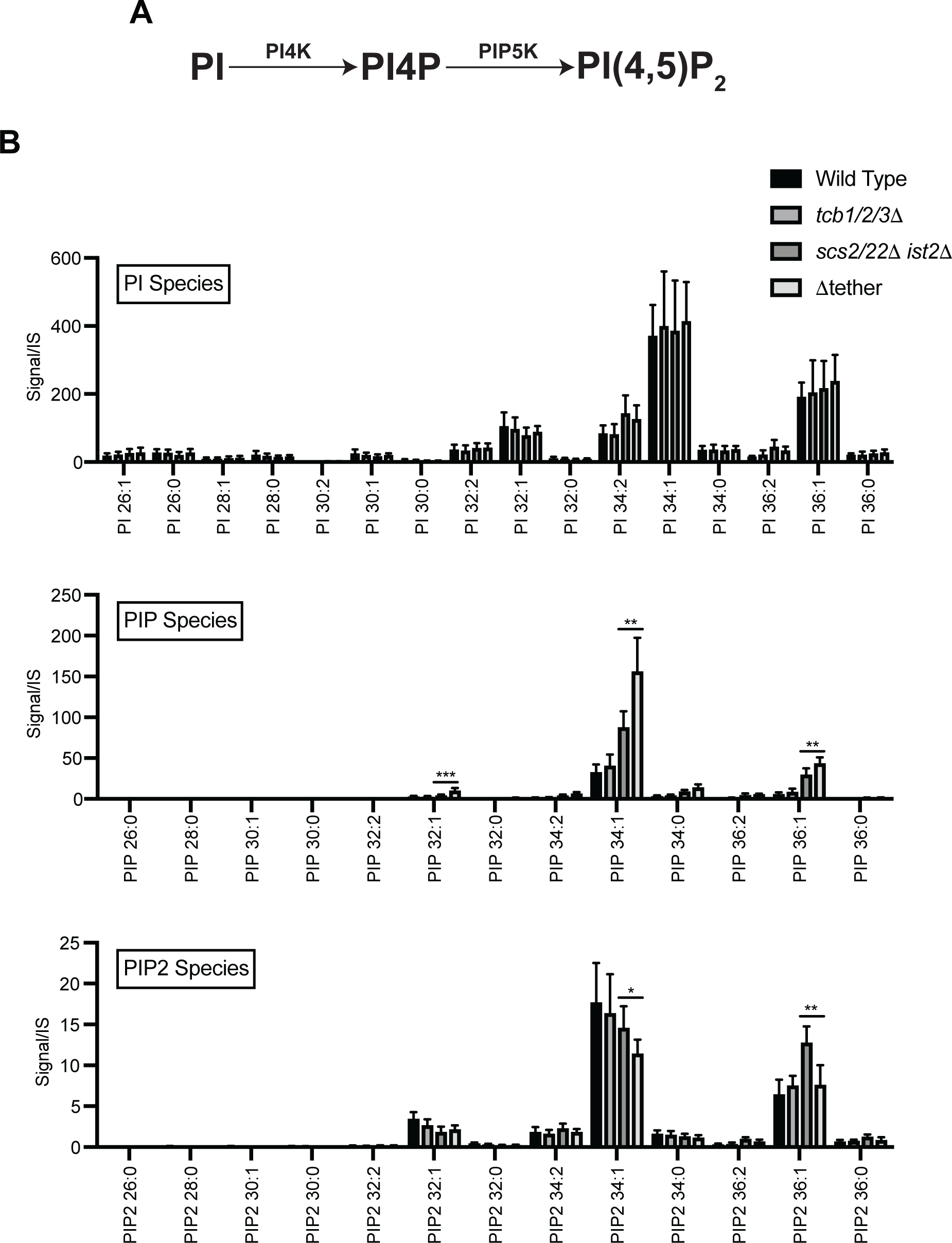
Lipidomic analyses of individual phosphoinositide species in the various ER-PM tether mutants. (**A**) Schematic representation of PI4P and PI(4,5)_2_ production from PI and the kinases involved. (**B**) Levels of PI, PIP, and PIP_2_ species in wild type, *tcb1/2/3*Δ, *scs2/22*Δ *ist2*Δ, and Δtether cells. Data represents mean ± standard deviation (N=5). * p > 0.1, ** p > 0.01.

**Figure S3.**
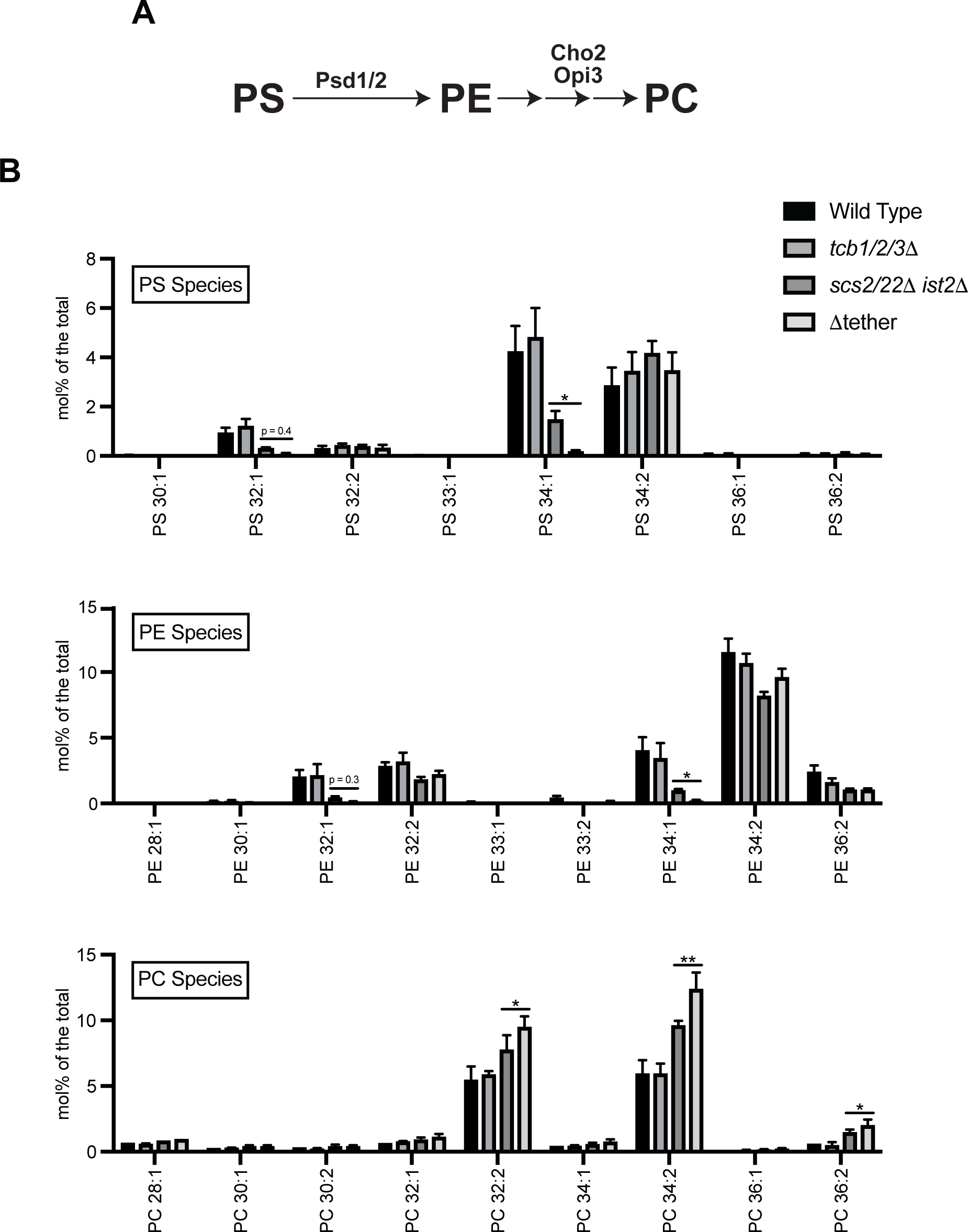
Lipidomic analyses of individual PS, PE, and PC species in the various ER-PM tether mutant strains. (**A**) Schematic representation of PE and PC production from PS and the enzymes involved. (**B**) Levels of PS, PE, and PC species in wild type, *tcb1/2/3*Δ, *scs2/22*Δ *ist2*Δ, and Δtether cells. Data represents mean ± standard deviation (N=3). * p > 0.1, ** p > 0.01.

**Figure S3.**
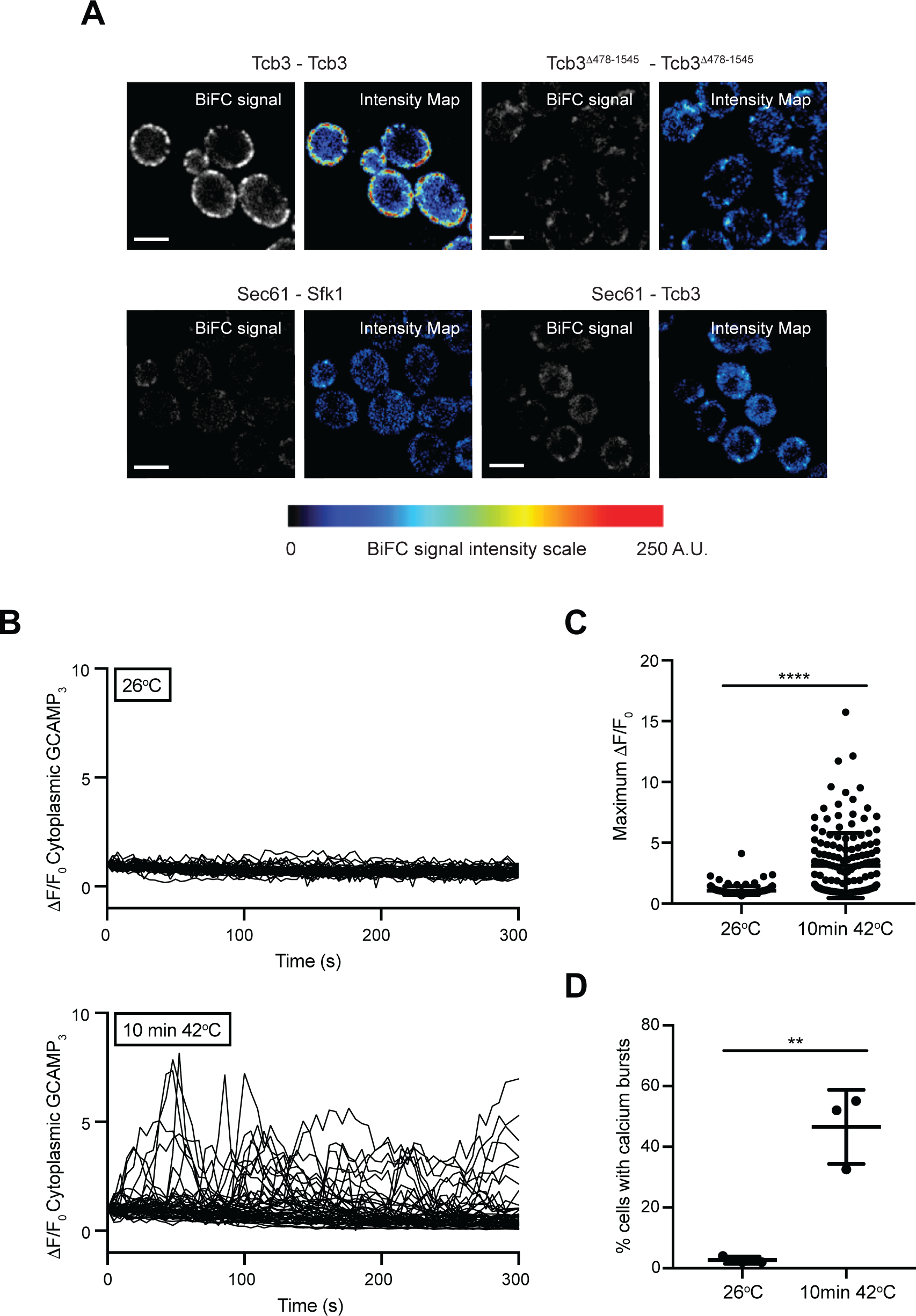
Tcb3 and Sfk1 display specifically associate, and heat stress results in cytoplasmic Ca^2+^ bursts. (**A**) Control protein-protein proximity assays; Protein-protein interactions between Tcb3 or Tcb3 ^Δ478-1545^ alone (top) or ER-localized Sec61 and either Tcb3 or Sfk1 (bottom) as detected by the split GFP BiFC assay. In each case, GFP_N_ is fused to the protein listed on the left and GFP_C_ is fused to the protein on listed on the right. (**B**) Normalised GCaMP3 fluorescence (ΔF/F_0_) traces of wild type cells incubated at the indicated temperature. Each line represents an individual cell, 49 (top) and 48 (bottom) traces for each condition. (**C**) Mean maximum normalised fluorescence (ΔF/F_0_) in wild type cells incubated at the indicated temperatures. Data represents mean ± standard deviation. Total number of cells analysed in three independent experiments: wild type 26°C, n=150; wild type 10 min 42°C, n=151. **** p > 0.0001. (**D**) Quantitation of the percentage of cells undergoing a calcium burst within 5 minutes after incubation at the temperature indicated. Data represents mean ± standard deviation in three independent experiments (n=50 cells per experiment). **** p > 0.0001.

**Figure S5.**
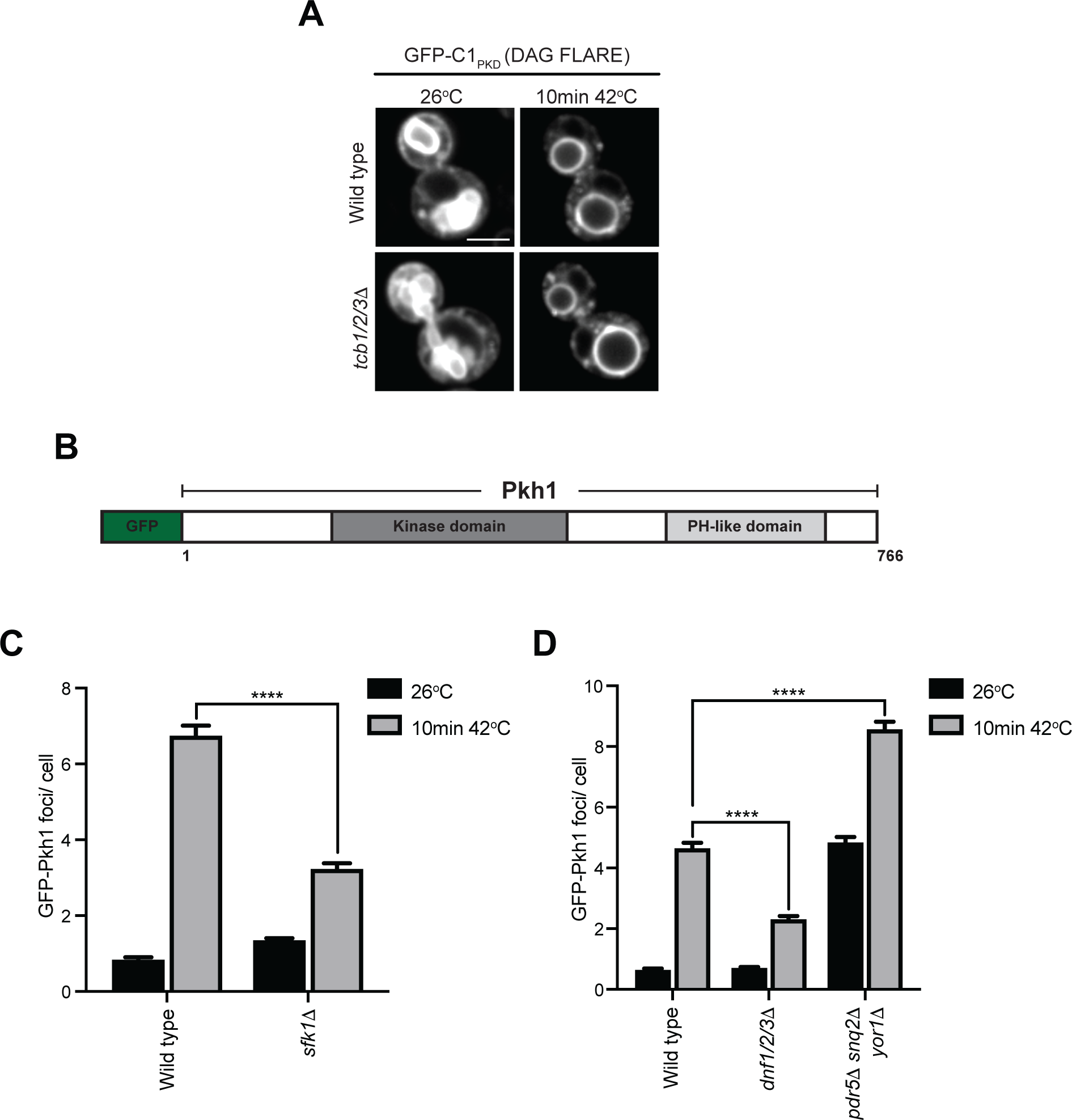
Heat-induced Pkh1 cortical puncta are regulated by Sfk1 and PM phospholipid asymmetry. (**A**) DAG FLARE (GFP-C1_PKD1_) localisation in wild type and *tcb1/2/3*Δ cells at 26°C or after a brief heat shock (10 min 42°C). Scale bar, 4 μm. (**B**) Schematic representation of GFP-Pkh1. (**C**) Quantitation of the number of GFP-Pkh1 puncta per cell in wild type and *sfk11*Δ cells at the indicated temperatures. Total number of cells analysed in three independent experiments: all strains and conditions, n=300). **** p > 0.0001. (**D**) Quantitation of the number of GFP-Pkh1 foci per cell in wild type, *dnf1/2/3*Δ, and *prd5*Δ*snq*2Δ*yor1*Δ cells at the indicated temperatures. Total number of cells analysed in three independent experiments: all strains and conditions n=400. **** p > 0.0001.

**Figure S6.**
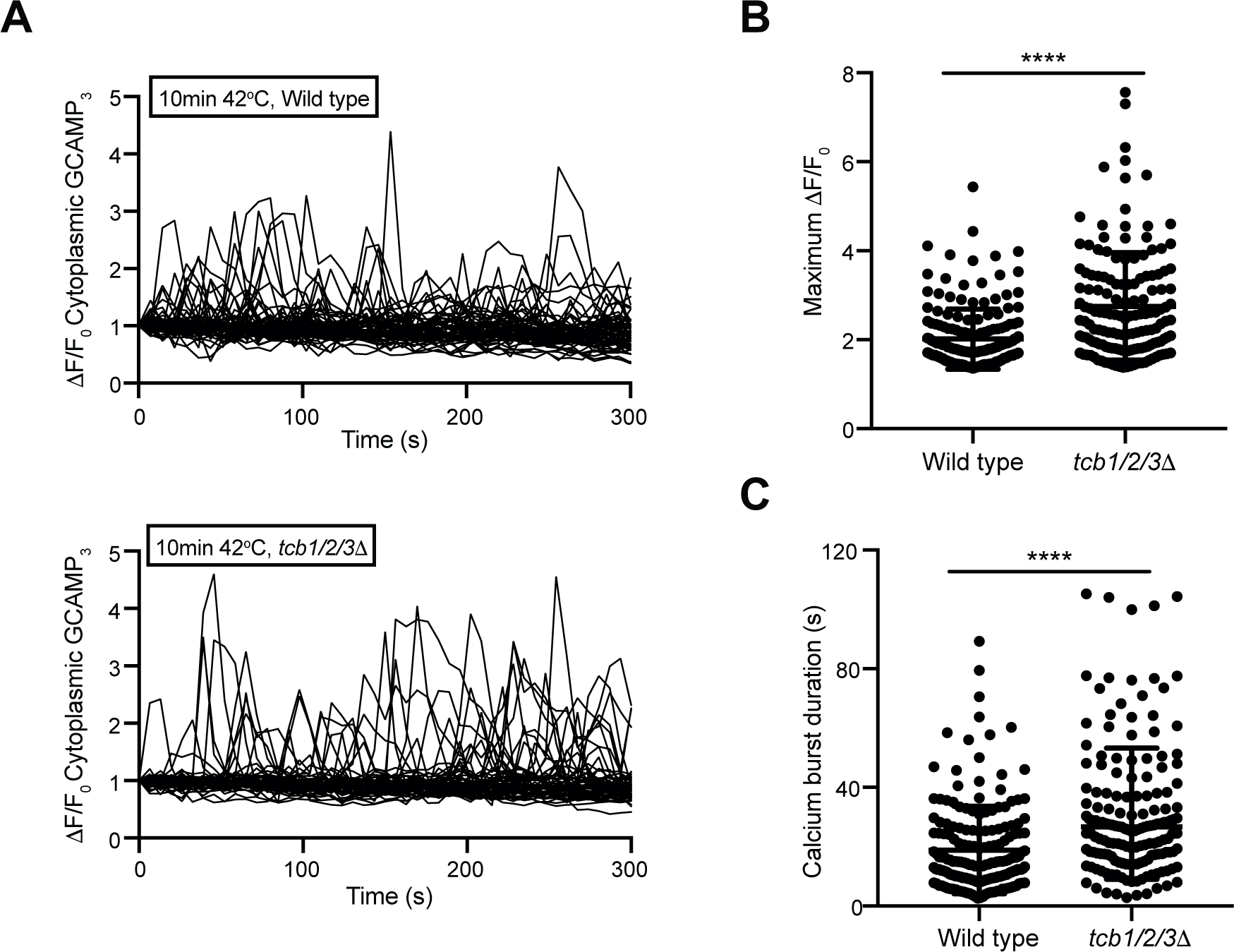
Tricalbin proteins regulate cytosolic Ca^2+^ burst intensity under heat stress conditions. (**A**) Normalised specific GCaMP3 fluorescence (ΔF/F_0_) traces of wild type and *tcb1/2/3*Δ cells incubated at 42°C for 10 min. Each line represents an individual cell, 62 (top) and 59 (bottom) traces for each condition. (**B** and **C**) Mean maximum normalized specific fluorescence (ΔF/F_0_) (**B**) and duration of calcium bursts (**C**) in wild type and *tcb1/2/3*Δ cells 5 minutes after incubation at the temperature indicated. Data represents mean ± standard deviation. Total number of cells analysed in three independent experiments: wild type 10 min 42°C n=150, *tcb1/2/3*Δ 10 min 42°C n=148). **** p > 0.0001.

### Supplemental Tables

**Supplemental Table 1.**
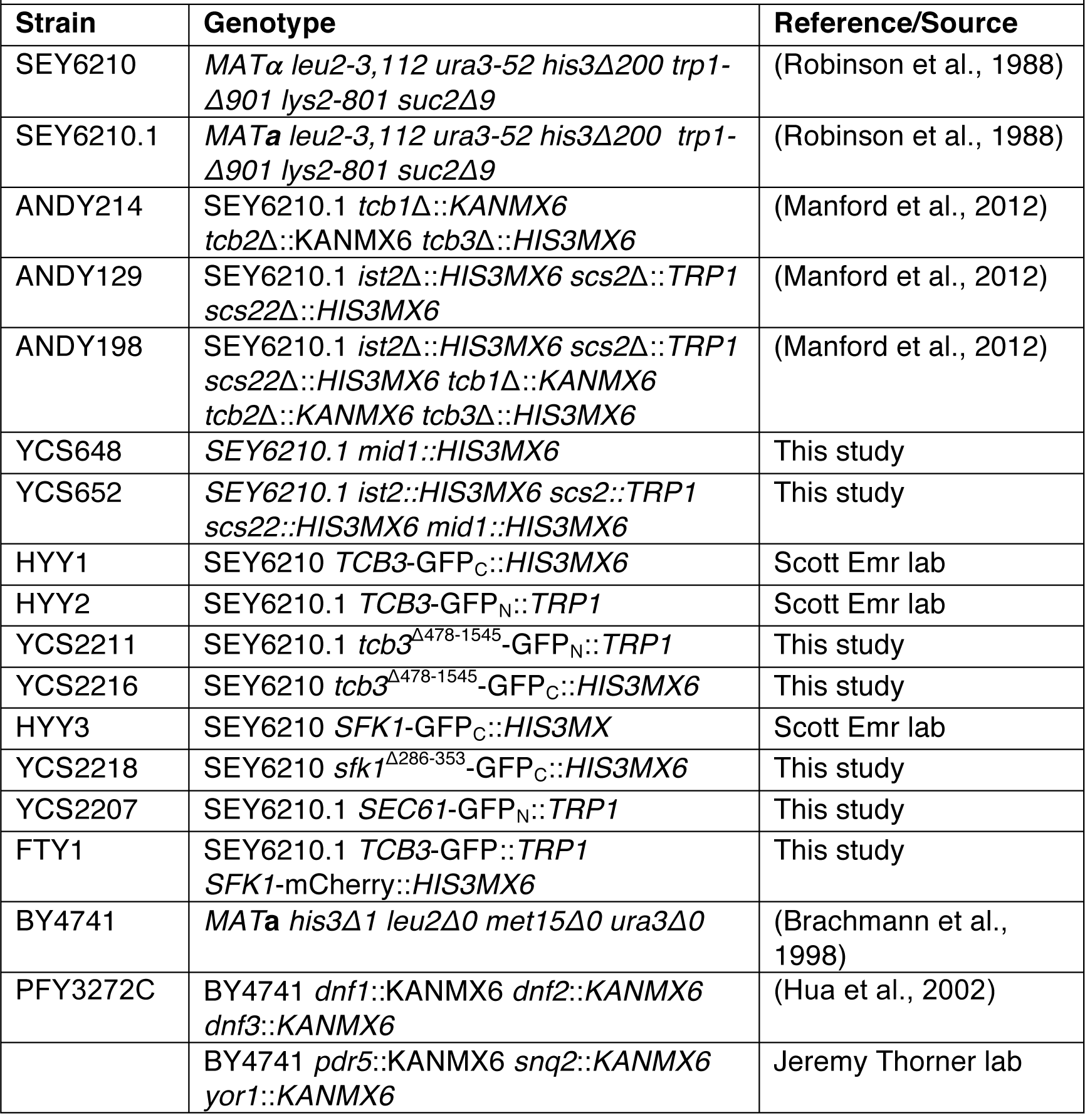
*Saccharomyces cerevisiae* strains used in this study.

**Supplemental Table 2.** Plasmids used in this study.

| **Plasmid Name** | **Description** | **Reference/Source** |
| --- | --- | --- |
|  | pRS416-p*GPD*-GFP-P4C<sub>SidC</sub> | (Luo et al., 2015) |
| pCS189 | pRS426-p*PRC1*-GFP-2xPH<sub>PLCδ</sub> | (Stefan et al., 2002) |
| pCS468 | pRS415-p*GPD*-GCaMP3 | (Omnus et al., 2016) |
|  | pRS416-p*GPD*-GFP-Lact-C2 | (Fairn et al., 2011) |
|  | pRS416-p*GPD*-mCherry-Lact-C2 | (Fairn et al., 2011) |
| pCS174 | pRS426-GFP-C1<sub>PKD</sub> | This study |
| pCS459 | pRS416-p*PKH1*-GFP-*PKH1* | (Omnus et al., 2016) |
| pAM43 | pRS415-p*TCB3-TCB3*-GFP-t*ADH1* | (Manford et al., 2012) |
| pAM44 | pRS415-p*TCB3-tcb3*<sup>Δ478-1545</sup>-GFP-t*ADH1* | (Manford et al., 2012) |
| pCB2 | pRS415-p*TCB3-tcb3*<sup>ΔSMP</sup>-GFP-t*ADH1* | (Collado et al., 2019) |
| pAM38 | pRS415-p*ADH1*-GFP-MSP<sub>Scs2</sub>-*SAC1* | (Manford et al., 2012) |

